# Integrative structure determination of a human mitochondrial contact site and cristae organizing system (MICOS) sub-assembly

**DOI:** 10.64898/2026.07.19.739404

**Authors:** Muskaan Jindal, Rakesh Mahato, Sreemoyee Das, Arko Guha, Kartik Majila, Shreyas Arvindekar, Anand T. Vaidya, Shruthi Viswanath

**Author notes:** Correspondence (A.T. V.) and (S.V).

## Abstract

The **Mi**tochondrial contact site and **C**ristae **O**rganizing **S**ystem (MICOS) complex is an inner mitochondrial membrane (IMM) assembly present at the cristae junction. It is responsible for regulating cristae formation and remodeling. However, its structure is not known. We applied Bayesian integrative structure determination to characterize the structure of the Mic60, Mic19, Mic10, and Mic13-containing MICOS sub-assembly combining AlphaFold predictions with data from crosslinking mass spectrometry, biochemical assays, electron tomography, homology modeling, and sequence alignments. The integrative structure revealed novel mutual interfaces among Mic10^N,C^, Mic60^LBS1,LBS2,mitofilin^, and Mic13^central,C^. Several likely-pathogenic missense mutations also localize to these novel interfaces, highlighting their importance. Further, we investigated some of these interactions using binding assays on purified proteins and demonstrated direct interactions among Mic10-Mic13-Mic60. Our results support a bridging role for Mic13 in the MICOS assembly, by binding Mic10 in the IMM-proximal region and Mic60 in the intermembrane space. Taken together, our integrative approach sheds light on the structure and assembly of the MICOS complex.

## Introduction

The inner mitochondrial membrane (IMM) contains the respiratory chain complexes, together with the mitochondrial contact site and cristae organizing system (MICOS) complex [1], [2], [3]. The MICOS complex is primarily present at the cristae junction (CJ). Along with ATP synthase and OPA1 GTPase, MICOS plays a critical role in forming and remodeling cristae [4], [5], [6]. The absence of some MICOS subunits results in alteration of cristae morphology or complete loss of cristae junctions [6], [7], [8], [9]. Mutations in several MICOS subunits have been associated with mitochondrial encephalopathy with liver dysfunction, mitochondrial myopathy with lactic acidosis, Parkinson’s disease, and type 2 diabetes mellitus [10], [11], [12].

The human MICOS complex contains seven subunits, organized as two sub-complexes that can exist independently: a Mic60 sub-complex, composed of Mic60, Mic19, and Mic25 (a paralog of Mic19), and a Mic10 sub-complex, composed of Mic10, Mic13, Mic26, and Mic27 (Fig. 1) [13]. Most of these subunits are conserved from yeast to humans [14]. Mic60, Mic10, Mic13, Mic26 and Mic27 are transmembrane proteins in the IMM, whereas Mic19 and Mic25 are peripheral membrane proteins in the intermembrane space (IMS) (Table S1) [1]. Owing to their membrane-bending properties, Mic60 and Mic10 are postulated to play a role in defining the curvature of cristae [15], [16], [17], [18]. Mic19 promotes Mic60 oligomerization and membrane remodeling activity [17]. Mic13 is known to play a role in stabilizing Mic10 [19], [20]. Mic26 and Mic27 are known to regulate Mic10 oligomerization in a cardiolipin-dependent manner [21].

**Figure 1.**
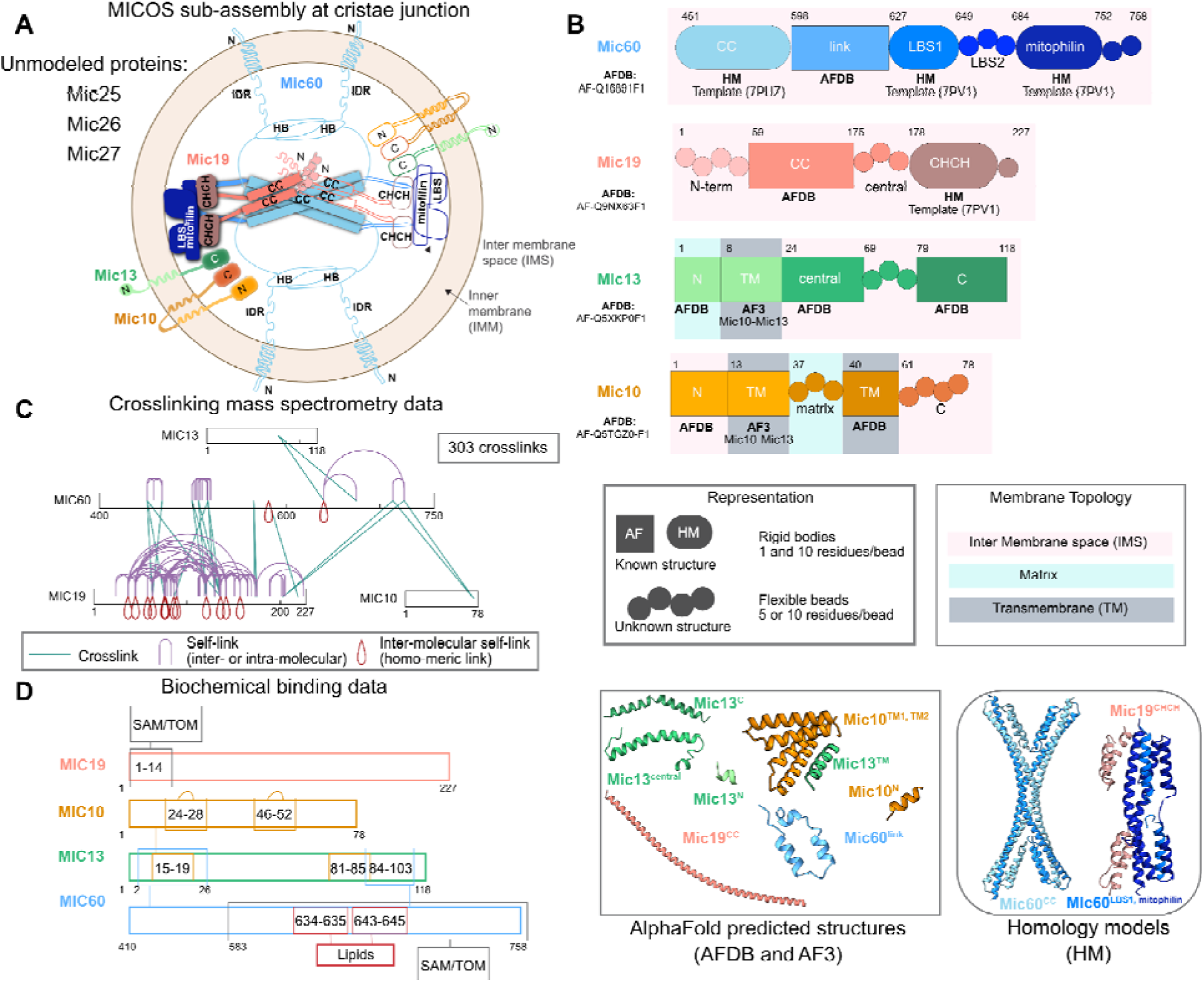
Information used in the modeling of the MICOS sub-assembly. **(A)** Schematic illustrating the components of the MICOS complex in the context of the holo-complex from a top view of the cristae junction. Th modeled MICOS protein domains are shaded in gradients from lighter to darker tones, from the N-terminal to the C-terminal end. Proteins with solid-colored regions indicate modeled regions, whereas outlined regions indicate regions that were not modeled. Mic60 and Mic19 are shown to be positioned at the CJ (shaded borders), whereas the other proteins are positioned below them, extending further into the cristae. (**B)** Regions with known atomic structure from AlphaFold and homology modeling are represented by rectangles and squircles, respectively, whereas the regions with unknown structure are represented by circles. The membrane topology is indicated by the background colors. **(C)** Crosslinks used in the study are depicted using xiNET [27]. **(D)** Protein-protein interaction data from biochemical studies are depicted by rectangles of the same color [15], [16], [28], [29], [30]. See also Tables S1, S3, S5. Abbreviations: N: N-terminal, C: C-terminal, TM: Transmembrane, CC: Coiled-coil, LBS: Lipid binding site, CHCH: Coiled-coil-helix-coiled-coil-helix, HB: Helical Bundle, IDR: Intrinsically Disordered Region, AFDB: AlphaFold database, AF3: AlphaFold3, HM: Homology Model, SAM: Sorting and Assembly Machinery, and TOM: Translocase of the Outer Membrane.

Despite its importance, the structure of the MICOS complex is not yet reported. Structures for parts of the MICOS complex, including a Mic60 coiled-coil (Mic60^CC^) tetramer and a 2:2 Mic60^LBS1,mitofilin^:Mic19^CHCH^ hetero-tetramer, have been characterized by X-ray crystallography in *Lachancea thermotolerans* and *Chaetomium thermophilum*, respectively (Fig. 1) [5], [22]. Recently, the structure of the Mic60^198-378^ helical bundle (HB) domain has been resolved using X-ray crystallography, and the study has shed light on the dynamics of the Mic60-Mic19 subcomplex and its role as a diffusion barrier [23]. However, no experimental structures exist for the remaining subunits or subcomplexes. Further, the mechanism by which the two MICOS sub-complexes are assembled into the MICOS complex is also not known.

Structures of large macromolecular assemblies such as the MICOS complex are often challenging to determine using a single experimental technique such as X-ray crystallography or cryo-electron microscopy. Integrative structure determination offers a useful alternative approach in such cases. In this approach, information from experiments is combined with physical principles, statistics of known structures, and prior models for structure determination [24], [25], [26]. Bayesian integrative structure determination *via* the open-source Integrative Modeling Platform (IMP; https://integrativemodeling.org) allowed us to combine sparse, noisy, ambiguous, low-resolution data from multiple sources at different spatial resolutions in an unbiased manner [24], [25], [26].

Here, we obtained the integrative structure of the human 4:2:2:1 Mic60^IMS^:Mic19:Mic10:Mic13 MICOS sub-assembly; the structure contains full-length Mic19, Mic10, and Mic13, and the IMS region of Mic60 (Mic60^451-758^). AlphaFold (AF) predictions allowed us to increase the structural coverage (percentage of residues with known experimental structure or homology model) by 55%, on average, per subunit. Complementarily, hundreds of *in vivo* and *in vitro* chemical crosslinks from crosslinking mass spectrometry (XLMS) are available. We combined information from AF predictions, XLMS, biochemical assays, electron tomography (ET), homology modeling, and sequence alignments to obtain the structure of the MICOS sub-assembly at a precision of 16 Å.

Refining previously known interactions, the integrative structure revealed novel mutual interfaces among Mic60^LBS1,LBS2,mitofilin^, Mic10^N,C^, and Mic13^central,C^. Several missense, likely-pathogenic mutations map to these novel interfaces, highlighting their importance. These interfaces guided subsequent *in vitro* binding experiments in *C. thermophilum*, which demonstrated direct interactions between Mic60^IMS^ and Mic13 and between Mic10^C^ and Mic13. Our experiments and integrative structural modeling indicate that Mic13 likely facilitates MICOS assembly by binding to Mic10 near the IMM *via* its transmembrane and central domains, and to Mic60 in the IMS *via* its central and C-terminal domains. Taken together, our integrative structural biology approach, combining modeling and experiments, reveals the interactions between Mic10 and Mic60 sub-complexes that likely maintain the structural integrity of the MICOS complex.

## Results

### AlphaFold predictions

First, we used AlphaFold3 (AF3) to predict the structures of the MICOS sub-assembly, composed of Mic10, Mic13, Mic60, and Mic19 (Table S2, Methods). However, the AF3 predictions of the larger 4:2:2:1 Mic60^IMS^:Mic19:Mic10:Mic13 MICOS sub-assembly as well as the 1:1:1:1 Mic60^IMS^:Mic19:Mic10:Mic13 sub-assembly were not confident. Therefore, we used AF3 to predict structures of various smaller sub-complexes of MICOS subunits. Among the homo-oligomers of the MICOS subunits, only the Mic10 homodimer prediction was confident. Next, AF3 was used to predict structures of hetero-oligomers of the MICOS subunits. The AF3 prediction of 1:1 Mic10-Mic13 was confident, but this prediction was inconsistent with the known membrane topology of these proteins: *e.g.*, the Mic13^IMS^ domains were positioned alongside the Mic10^TM^ domains. However, the prediction on a truncated transmembrane version of the two proteins, 2:1 Mic10^TM1,matrix,TM2^:Mic13^TM^, was confident and consistent with the confident Mic10 homodimer prediction reported above. This confident 2:1 Mic10^TM1,matrix,TM2^:Mic13^TM^ prediction, along with additional confidently predicted domains from monomeric AlphaFold predictions of MICOS subunits, was used to define rigid bodies for integrative modeling (Table S1, S2). Further, the AF3 predictions of 1:1 Mic60^LBS1,LBS2,mitofilin^:Mic10 and 1:1 Mic60^LBS1,LBS2,mitofilin^:Mic13^central,C^ were also confident, but the Mic60 structures in these predictions were inconsistent with the reported crystal structure of Mic60^LBS1,mitofilin^ (RMSD = 17 Å). Therefore, instead of using these predictions to define rigid bodies, we used the high-probability interface residue pairs from these predictions to define distance restraints between the corresponding Mic60-Mic10 and Mic60-Mic13 residues (Table S3, Methods). Overall, AF predictions allowed us to increase the structural coverage by 72%, 91%, 53%, and 6% for Mic10, Mic13, Mic19, and Mic60, respectively.

### Stoichiometry of the modeled MICOS sub-assembly

Mic10, Mic13, Mic60, and Mic19 are core components of the MICOS complex [5]. Based on the AF structure predictions and other considerations, we modeled a 4:2:2:1 Mic60^IMS^:Mic19:Mic10:Mic13 MICOS sub-assembly corresponding to the MICOS subunits in one half of the CJ, *i.e.,* a half-junction of MICOS (solid-colored regions in Fig. 1, Table S1, Table S2). We also considered modeling the holo complex using symmetry (see Discussion). The stoichiometry of Mic60 and Mic19 was based on the X-ray structures of a Mic60^CC^ homo-tetramer, and a 2:2 Mic60^LBS1,mitofilin^:Mic19^CHCH^ hetero-tetramer [22]. The Mic60^CC^ tetramer is thought to reside in the center of the CJ, linking two Mic60^LBS1,mitofilin^:Mic19^CHCH^ heterotetramers anchored on opposite sides of the CJ [22]. Here, we modeled a MICOS sub-assembly occupying one half of the CJ, comprising one Mic60^CC^ tetramer and an associated Mic60^LBS1,mitofilin^-Mic19^CHCH^ heterotetramer, resulting in a 4:2 stoichiometric ratio of Mic60:Mic19. Mic10 exists as an oligomer in the MICOS complex [1], [15], [16]. In the absence of experimental structures, AF predictions of several Mic10 oligomers were obtained; only the Mic10 dimer prediction was confident, as noted above (Table S2). The stoichiometry of Mic13 in MICOS is unknown; given the lack of data, *e.g.*, crosslinks on Mic13, a single copy of Mic13 was modeled to minimize assumptions. We also attempted modeling with 2 copies of Mic13; however, AF3 predictions of 2:2 Mic10:Mic13 and 2:2 Mic60:Mic13 were not confident and could not be used as starting structures for the integrative modeling. Therefore, based on the above considerations and a confident 2:1 Mic10^TM^:Mic13^ΤΜ^ AF prediction, we modeled two copies of Mic10 and one copy of Mic13 (Table S2, Results: AlphaFold predictions). Finally, we did not include the N-terminus of Mic60, Mic60^1-450^, containing the transmembrane region of Mic60, Mic60^IDR^ and Mic60^HB^, Mic25 (a paralog of Mic19), Mic26, and Mic27 in our model, due to a lack of XLMS and other biochemical interaction data on these regions.

### Rationale for modeling the human MICOS complex

We chose to model the human MICOS sub-assembly given the greater availability of data from experimental studies and more confident AF predictions on the human complex. We also considered modeling the sub-assembly in *C. thermophilum*; however, the AF3 predictions for the Mic10-Mic13, Mic10-Mic60, and Mic13-Mic60 complexes in this case are not confident.

Experimental data from XLMS (46% of crosslinks), biochemical assays, and ET were derived from studies using human cells (Table S3-S5). The remaining crosslinks (54%) are from mouse cells, but these sequences, on average, have a high sequence similarity (>95%) to human sequences (Mic60: 97%, Mic19: 95%, Mic10: 96%, and Mic13: 94%), indicating that these mouse-derived crosslinks can be mapped to human sequences (Table S4). Finally, the homology models are based on template experimental structures in *C. thermophilum* and *L. thermotolerans*. Recently, other groups [23] have also used these structures to model the corresponding human Mic60 tetramer and Mic60-Mic19 complexes. These modeled regions correspond to conserved regions, based on ConSurf scores and the percentage of residues with 70% sequence conservation, based on alignments in prior studies (Fig. S3) [22]. Although the sequence similarity between the modeled human regions and the species corresponding to the experimental structure is ∼43%, these regions are known to form coiled-coils and helices, with the residues in the interface and helix-break regions conserved (Fig. S3). These homology models typically achieve ∼3-5 Å RMSD accuracy, with greater uncertainty in side-chain conformations and loop regions [31]. However, this degree of accuracy suffices for the coarse-grained modeling presented here, which uses representations of 1 and 10 residues per bead.

### Summary of the integrative modeling workflow

The integrative modeling was performed in four stages (Fig. 2, Methods). Information from XLMS, biochemical assays, and electron tomography (ET) was integrated with AF predictions, homology models, sequence alignments, and stereochemistry information (Fig. 1, 2). Each protein was represented as a string of spherical beads. For computational efficiency, the regions with known atomic structures were represented simultaneously at 1 and 10 residues per bead and modeled as rigid bodies; in a rigid body, the relative positions and orientations between the beads are fixed (Table S1). Whereas, the regions with unknown structure were represented at 5 or 10 residues per bead and modeled as flexible beads which can move relative to one another (Fig. 1, Table S1, Methods). Data from XLMS was used to restrain the distances between beads corresponding to the crosslinked residues (Table S4). Similarly, interactions from co-IP and AF-predicted interfaces were used to restrain the distances between beads corresponding to the interacting domains and interface residue pairs, respectively (Table S3). The membrane thickness from ET, together with protein subcellular fractionation and liposome sedimentation data, was used to localize protein domains according to their membrane topology (Fig. S1, Table S5, Methods).

**Figure 2.**
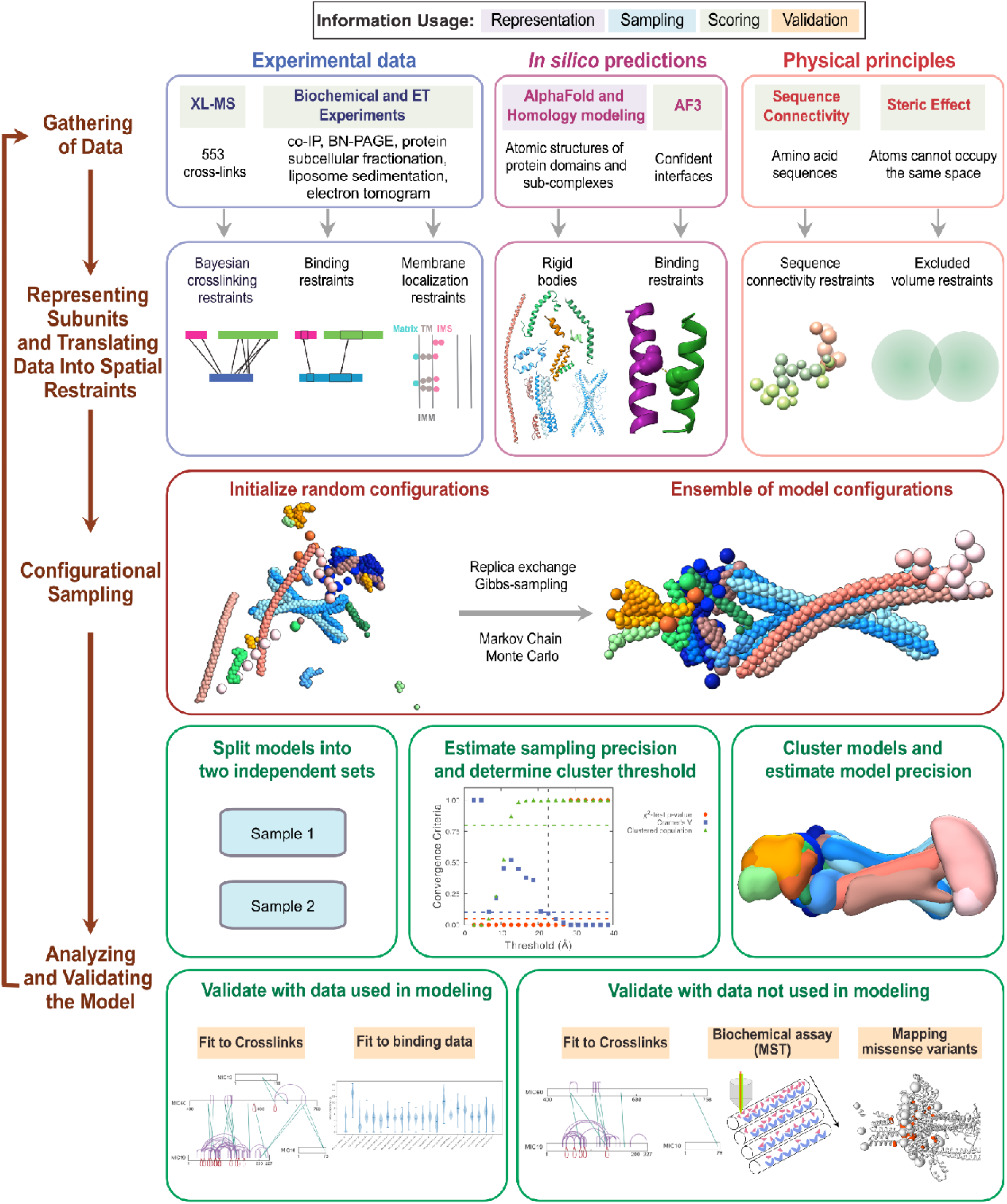
Integrative structure determination of the MICOS sub-assembly. Schematic of the four stages of th integrative modeling pipeline. The first row describes the input information, the second row shows how each input information is used to encode restraints, the third row depicts the sampling method, and the last two rows illustrate the analysis and validation steps used in the modeling. The background colors in the input information correspond t the stage in which the information is used, as mentioned in the legend at the top. Abbreviations: co-IP: co-Immunoprecipitation, BN-PAGE: Blue native polyacrylamide gel electrophoresis, and MST: Microscale Thermophoresis.

We used the Gibbs sampling Replica Exchange Markov Chain Monte Carlo algorithm. At each Monte Carlo step, the models are scored using restraints derived from the input information (above) along with stereochemistry restraints such as sequence connectivity and excluded volume. We sampled ∼200 million models from 50 independent runs, with each run starting from a random initial configuration. The analysis and validation protocol were similar to those in previous studies (Fig. 2, Fig. S2, Methods) [32], [33], [34], [35], [36], [37].

### Integrative structure of the MICOS sub-assembly

Integrative modeling of the 4:2:2:1 Mic60:Mic19:Mic10:Mic13 sub-assembly resulted in a single major cluster of 27,832 models (93% of 30,000 models), with a model precision of 16 Å (Fig. S1, Methods). Model precision is the variability of models in this cluster, defined by the average RMSD of the cluster models to the cluster centroid. The models fit well with the input crosslinks, biochemical binding data, membrane localization data, and AF-predicted contacts used in modeling (Fig. S4-S5, Table S6). The models also fit well with the crosslinks not used in modeling; the few violations correspond to crosslinks on Mic19^N, CC^ whose structure is not known (Fig. S6, Table S6, see Discussion).

The resulting integrative structure was visualized in two ways: a representative bead model corresponding to the centroid of the major cluster (Fig. 3A), and localization probability density maps, representing the localization of protein domains (Fig. 3B). These maps specify the probability of a voxel (3D volume element) being occupied by a domain in the set of structurally superposed cluster models.

**Figure 3.**
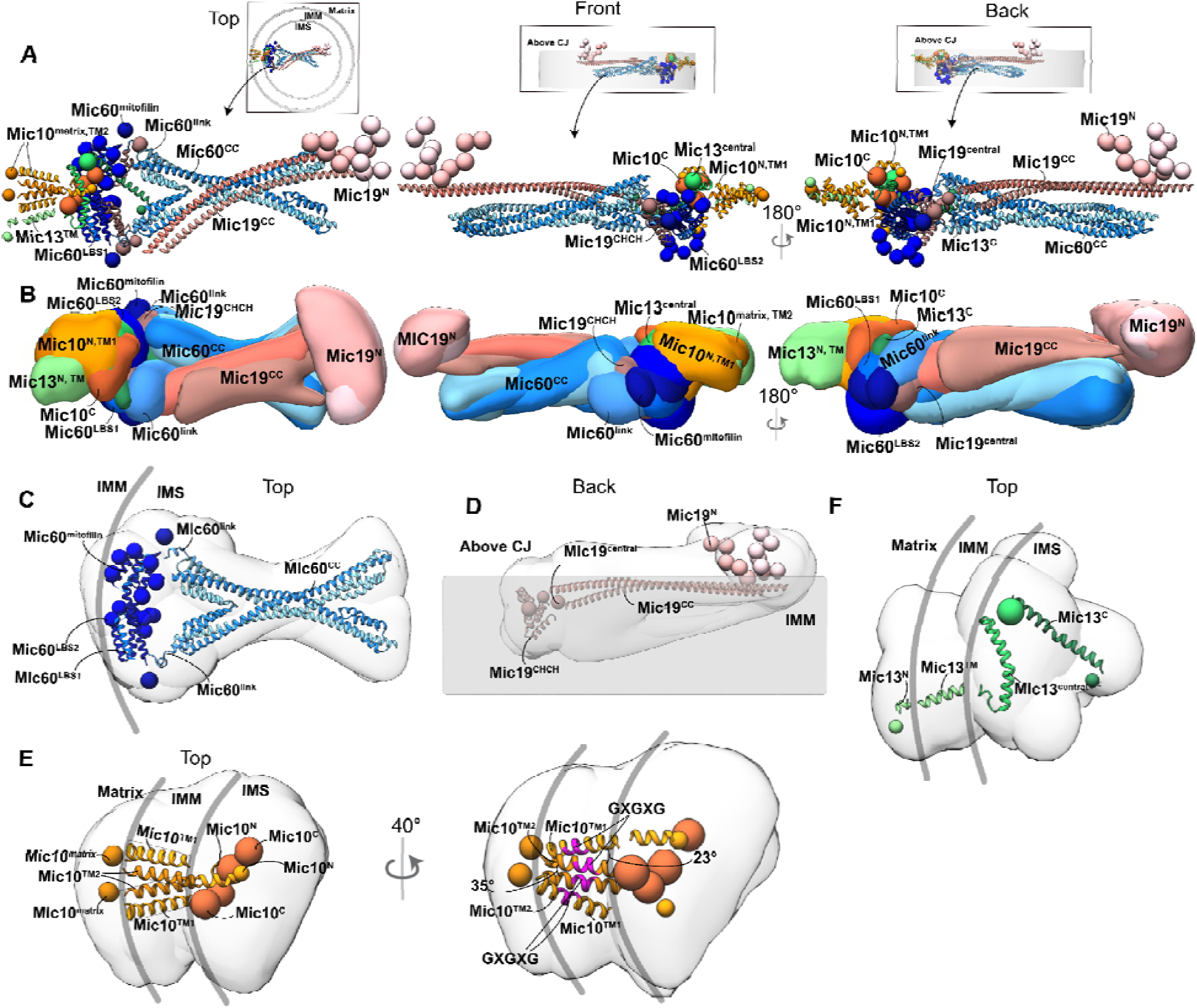
Integrative model of the MICOS sub-assembly. **(A)** Representative cluster center bead model of the most populated cluster of integrative models of the 4:2:1:1 Mic60:Mic19:Mic10:Mic13 MICOS sub-assembly. Regions with known atomic structures are represented as ribbons, and the regions with unknown structure are depicted as spherical beads. The proteins are shaded in gradients from lighter to darker tones, from the N-terminal to the C-terminal end. Domains of Mic60 (blue), Mic19 (salmon), Mic13 (green), and Mic10 (orange) are marked. Insets indicate the membrane orientation of each view. **(B)** Localization probability densities show the location of eac domain in the cluster. The threshold for each density is set to ∼10% (except for Mic19^central^:4%, Mic13^N^:26%, MicMic19^CC^, and Mic19^N^: 30%). **(C-F)** The domains of each modeled MICOS subunit in the bead model are shown; their corresponding densities are shown in the background in light gray. The boxes in the top figure of panel **E** delineate the two copies of Mic10, and the GXGXG motifs are colored in magenta in the bottom figure.

The modeled MICOS sub-assembly extends into the CJ, with a protein-dense region proximal to the IMM. Mic10 and Mic13 anchor the sub-assembly at the membrane, interacting with the Mic60^LBS1,LBS2,mitofilin^-Mic19^CHCH^ hetero-tetramer to form the membrane-proximal protein-dense region. Additionally, the Mic60 and Mic19 coiled-coil domains extend into the CJ. Next, we describe the localization and domain organization of each MICOS subunit.

#### Mic60

The domains of Mic60 appear to form a triangular architecture at the protein-dense region, with Mic60^CC^ constituting two sides of the triangle and Mic60^LBS1-LBS2-mitofilin^ forming the other side (Fig. 1, Fig. 3C, Table S1, Table S7). Mic60^CC^ is oriented in the plane of the CJ. The Mic60^LBS1-LBS2-mitofilin^ side of the triangle is flanked on both sides by the short Mic60^link^ domain that connects the Mic60^CC^ to Mic60^LBS1^. All the domains of Mic60, except for parts of Mic60^CC^ and Mic60^LBS2^ are annotated as high-precision regions (Fig. S7).

#### Mic19

The two copies of Mic19^CC^, modeled as independent helices from AF, form a parallel dimer extending in the IMS across the CJ (Fig. 1, Fig. 3D, Table S1, Table S7). Mic19^N^, a region of unknown structure, localizes above the CJ and is likely to interact with the SAM complex [38], [39]. Mic19^CHCH^, complexed with Mic60^LBS1-mitofilin^, is anchored near the membrane. The Mic19^CHCH^ is annotated as a high-precision region, whereas the Mic19^N^ and parts of the Mic19^CC^ domains are low-precision regions (Fig. S7).

#### Mic10

In the AF-predicted 2:1 Mic10^TM^-Mic13^TM^ structure, each Mic10 monomer forms a wedge-shaped architecture, with the two transmembrane helices of each Mic10 monomer oriented at approximately 23° relative to each other [15], [16] (Fig. 1, Fig. 3E, Table S1, Table S7). Additionally, in the integrative structure, the Mic10^N^ and Mic10^C^ of a Mic10 monomer are proximal in the IMS (Fig. 3E, Table S7). Further, the two Mic10 monomers in the AF-predicted structure are oriented such that their Mic10^TM2^ helices are at approximately 35° relative to one another (Fig. 3E). In the integrative structure, the Mic10 monomers interact *via* both their Mic10^TM2^ (GXGXG motif) helices and Mic10^C^ domains (Fig. 3E, Table S7). Previously, it was shown that the GXGXG motifs in its transmembrane helices are involved in Mic10 oligomerization; here, we provide a structural basis for this observation [16]. It is possible that the repetition of this dimeric Mic10 arrangement, defined by these inter-monomer and intra-monomer angles, may collectively promote membrane curvature, providing a structural basis for membrane bending by Mic10. Regions of Mic10^N^, Mic10^TM1,TM2^, and Mic10^C^ in the Mic10 monomer that is proximal to Mic13 are of high-precision and the region of Mic10^N^ in the other Mic10 monomer is of low-precision (Fig. S7).

#### Mic13

Mic13 comprises Mic13^N^, an N-terminal helix region anchored in the matrix; Mic13^TM^, a transmembrane helix; and two regions in the IMS, Mic13^central^ and Mic13^C^, both of which are confidently predicted as helices in AF (Fig. 1, Table S2, see Methods). Mic13^central^ is an amphipathic helix, localized parallel to the IMM in most models (Fig. 3F, Table S1). Regions of Mic13^C^ are localized at low-precision (Fig. S7).

### Protein-protein interactions in MICOS

Below, we describe the novel protein-protein interfaces between MICOS subunits identified from the integrative structure. Protein-protein interfaces are denoted as Mic10^N^-Mic60^mitofilin^, for example, to convey that the N domain of Mic10 forms an interface with the mitofilin domain of Mic60.

#### Mic10-Mic13

In the AF-predicted 2:1 Mic10^TM^:Mic13^TM^ structure, Mic13^TM^ is positioned to one side of the complex, adjacent to one of the Mic10^TM^ helices. Previously, it was shown that the Mic10-Mic13 interaction involves the transmembrane GXXXG and C-terminal WN motifs of Mic13, but the corresponding interacting region on Mic10 has not been experimentally identified [30]. In this structure, we show that the GXXXG motif of Mic13 interacts with the GXGXG motif of Mic10^TM1^ and Mic13^WN^ interacts with Mic10^C^ (Fig. 1, Fig. 4A, Table S7). In addition, we identify a previously unreported Mic10^C^-Mic13^central^ interface.

**Figure 4.**
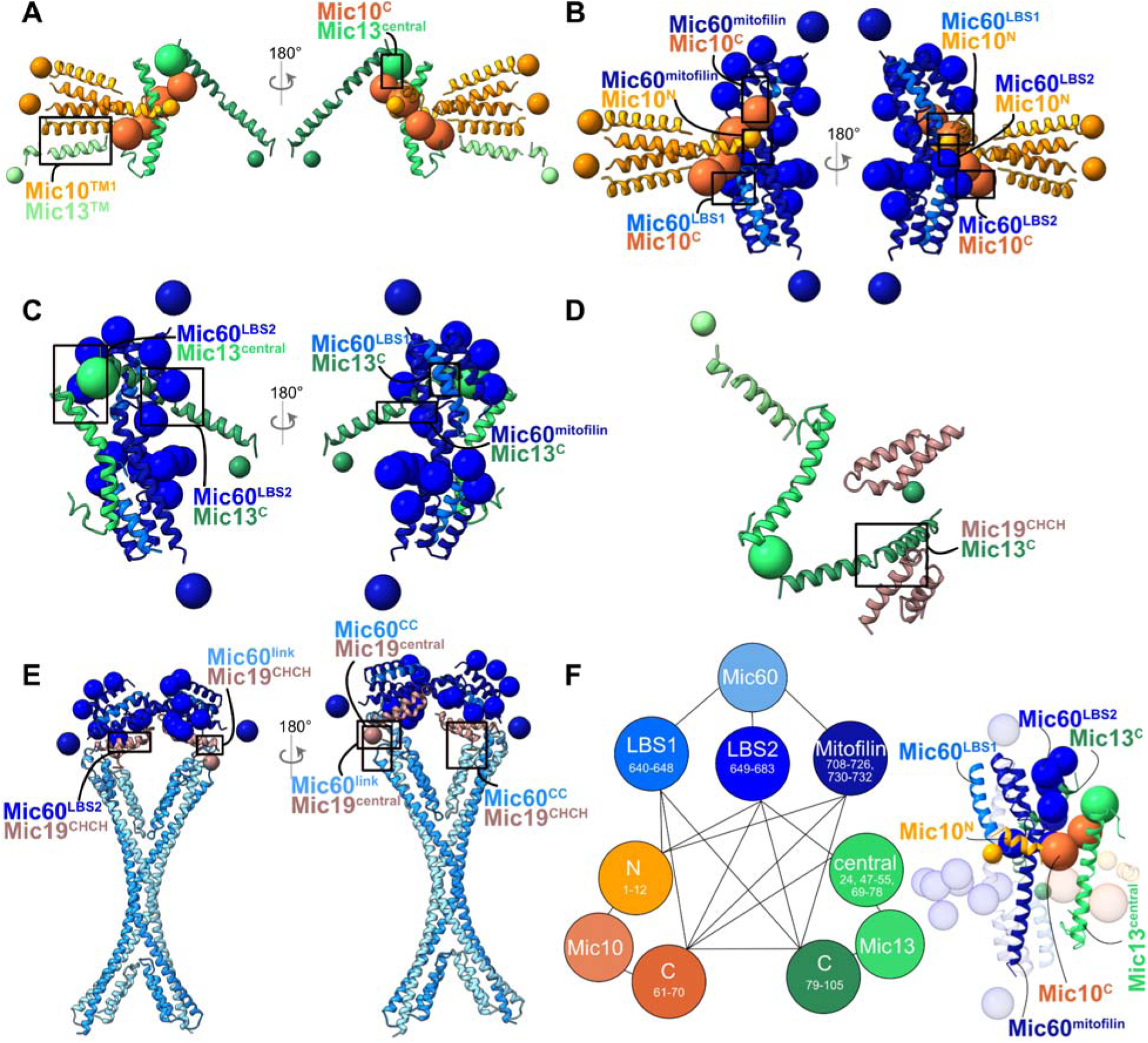
Interactions between the modeled MICOS protein domains. The protein-protein interfaces from the integrative structure are shown for **(A)** Mic10-Mic13, **(B)** Mic60-Mic10, **(C)** Mic60-Mic13, **(D)** Mic19-Mic13, and **(E)** Mic60-Mic19. The black boxes show the interfaces at the domain level. **(F)** Regions corresponding to mutual interactions among Mic10, Mic13, and Mic60 domains are shown *via* a network diagram (left) and on the bead model (right). M denotes the mitofilin domain in Mic60. For a detailed list of interfaces, see Table S7.

#### Mic60-Mic10

Although Mic60 and Mic10 are known to interact, their binding interface was previously undefined [16], [40], [41]. The integrative structure indicates that Mic60^LBS1,LBS2,mitofilin^ interacts with Mic10^N,C^ (Fig. 1, Fig. 4B, Table S7).

#### Mic60-Mic13

Mic60 is known to interact with Mic13^C^ (Mic13^84-103^); however, the corresponding interacting region in Mic60 has not been identified [30]. The integrative structure indicates that Mic60^LBS1,LBS2,mitofilin^ interact with Mic13^C^. In addition, we identified a novel Mic60^LBS2^-Mic13^central^ interface (Fig. 1, Fig. 4C, Table S7).

#### Mic19-Mic13

To our knowledge, no interaction between Mic19 and Mic13 has been reported so far. Our integrative structure reveals a Mic19^CHCH^-Mic13^C^ interface (Fig. 1, Fig. 4D, Table S7).

#### Mic60-Mic19

We also identified previously unknown interfaces between Mic60 and Mic19: Mic60^CC,link^-Mic19^central,CHCH^ and Mic60^LBS2^-Mic19^CHCH^, resulting from the triangular architecture of Mic60 domains (Fig. 1, Fig. 4E, Table S7).

#### Mutual interfaces among Mic10, Mic13, and Mic60

From the above-described interfaces, it is evident that there are mutual interfaces among Mic60^LBS1,LBS2,mitofilin^, Mic10^N,C^, and Mic13^central,C^. These mutual interfaces can be described in terms of these tripartite interfaces: Mic60^LBS1,LBS2^-Mic10^C^-Mic13^C^ (Mic60^648-683^-Mic10^61-70^-Mic13^89-92^), Mic60^mitofilin^-Mic10^C^-Mic13^C^ (Mic60^716-730^-Mic10^61-70^-Mic13^84-92^), and Mic60^LBS2^-Mic10^C^-Mic13^central^ (Mic60^664-668^-Mic10^61-70^-Mic13^24,47-55,69-78^) (Fig. 1, Fig. 4F, Table S7). Notably, the parts of the Mic13^central^, Mic10^C^, and Mic60^LBS2^ regions in these mutual interfaces correspond to regions of unknown structure (Fig. 1, Table S1). Further, Mic60^LBS1,LBS2,mitofilin^ and Mic10^C^ regions are also of high precision in the integrative structure (Fig. S7).

### Mic10-Mic13-Mic60 interactions using in vitro experiments

The interaction interfaces among Mic10, Mic13, and Mic60 identified from the integrative structural model guided subsequent *in vitro* characterization of purified MICOS subunits. Several attempts to purify different constructs of human and *Chaetomium thermophilum* (ct) homologs of Mic60, Mic13 and Mic10, have been unsuccessful. Hence, the ctMic60^IMS^ (residues 208-691), ctMic12 (*Chaetomium thermophilum* homolog of human Mic13), and synthetic peptides corresponding to ctMic10’s N-terminal (ctMic10^N^; residues 1–27) and C-terminal (ctMic10^C^; residues 76–93) regions were used to test protein-protein interactions *via* the Microscale Thermophoresis (MST) assay (Fig. 5, Fig. S8). The *Chaetomium thermophilum* homologs are more stable and have also been previously used for in vitro studies [17], [22]. MST measurements indicate an interaction between ctMic12 and ctMic10^C^ (K_d_ = 609 ± 175 nM), and between ctMic12 and ctMic60^IMS^ (K_d_ = 111 ± 24 nM, Fig. 5). No detectable binding was observed between ctMic12 and ctMic10^N^ or between ctMic60^IMS^ and the two peptides (ctMic10^C^, ctMic10^N^) (Fig. 5, Fig. S9). Also, no binding was observed between ctMic12 and 1xFLAG peptide, which acts as a negative control (Fig. S9).

**Figure 5.**
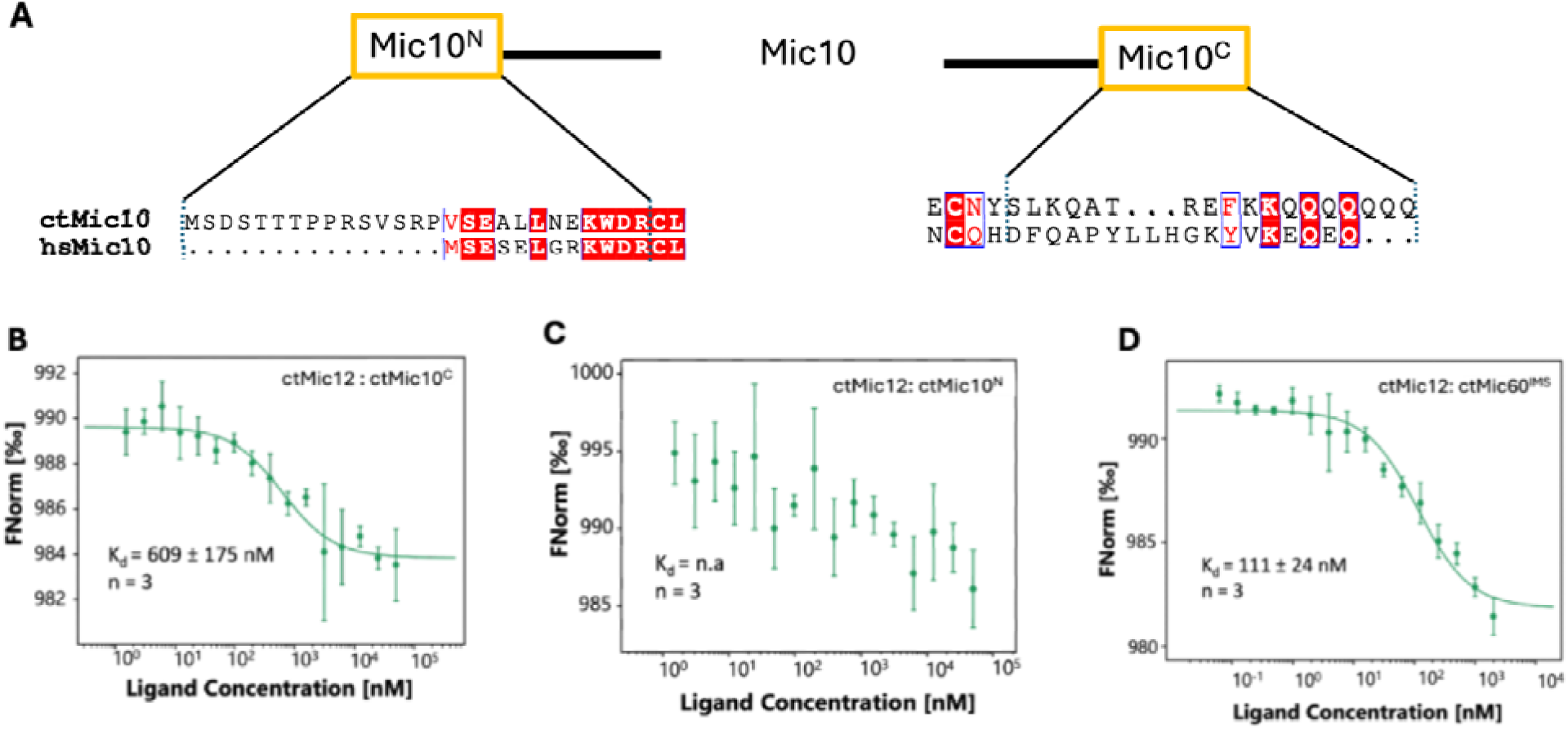
MST analysis of interactions between ctMic12, Mic10 peptides, and Mic60^IMS^. **(A)** The alignment of the N-terminal (Mic10^N^; left) and C-terminal (Mic10^C^; right) soluble regions of ctMic10 and hsMic10 show the domain-level conservation. The peptide ctMic10^N^ was terminated at the Arg (R) residue of the conserved KWDRCL patch, as the next Cys (C) residue is predicted to be part of the transmembrane helix. Conserved residues are in white, whil the similar amino acids are in red. **(B)** ctMic12 interacts with ctMic10^C^ with a K_d_ of 609 ± 175 nM. **(C)** No detectable interaction between ctMic12 and ctMic10^N^. **(D)** ctMic12 interacts with ctMic60^IMS^ with a K_d_ of 111±24 nM. In thes measurements, ctMic12 was fluorescently labeled and maintained at a constant concentration, while the partner protein/peptide was the ligand that was titrated. Normalized fluorescence (FNorm) is plotted as a function of ligand concentration. Error bars represent standard deviation from triplicate measurements.

The *in vitro* binding results, in combination with the interaction interfaces derived from the integrative structure, suggest that Mic13 may play a central role in MICOS assembly by bridging the Mic10- and Mic60-subcomplexes. Specifically, Mic13 appears to interact with Mic10 *via* its central and transmembrane regions, and with Mic60^IMS^ through its central and C-terminal domains, thereby facilitating the organization and assembly of the MICOS complex.

### Mapping ClinVar variants of uncertain significance

Next, we used the integrative structure to rationalize the effects of the missense mutations. We obtained 30 missense mutations of uncertain significance on the modeled MICOS subunits from ClinVar, of which 18 were predicted to be likely pathogenic by AlphaMissense [42], [43] (Table S8). Of these mutations, 12 localize to protein-protein interfaces in the integrative structure, with the majority of these interface mutations (7 of 12) occurring at the Mic10-Mic13-Mic60 mutual interfaces (Fig. 4, Fig. 6, Table S8). The other six interface mutations occur at the Mic19^CC^-Mic19^CC^, Mic19^CHCH^-Mic60^CC^, Mic19^CHCH^-Mic60^mitofilin^, Mic60^CC^-Mic60^mitofilin^, and Mic10^TM1^-Mic13™ interfaces, respectively. The remaining 6 of 18 mutations are localized in exposed domains, such as Mic60^CC^, Mic60^link^, Mic19^N^, and Mic19^CHCH^, which interact with the sorting and assembly machinery/topogenesis of the mitochondrial outer membrane β-barrel proteins (SAM/TOB) complex [8], [38], [39], [44]. Taken together, these mutations could be pathogenic because they may disrupt the protein-protein interfaces in the MICOS complex or in the larger MIB complex consisting of MICOS and SAM/TOB (Fig. 6). However, MICOS subunits have other mitochondrial functions, such as in mtDNA maintenance and in mitochondrial protein import, and the above MICOS subunit mutations may also be rationalized in these other contexts [45].

**Figure 6.**
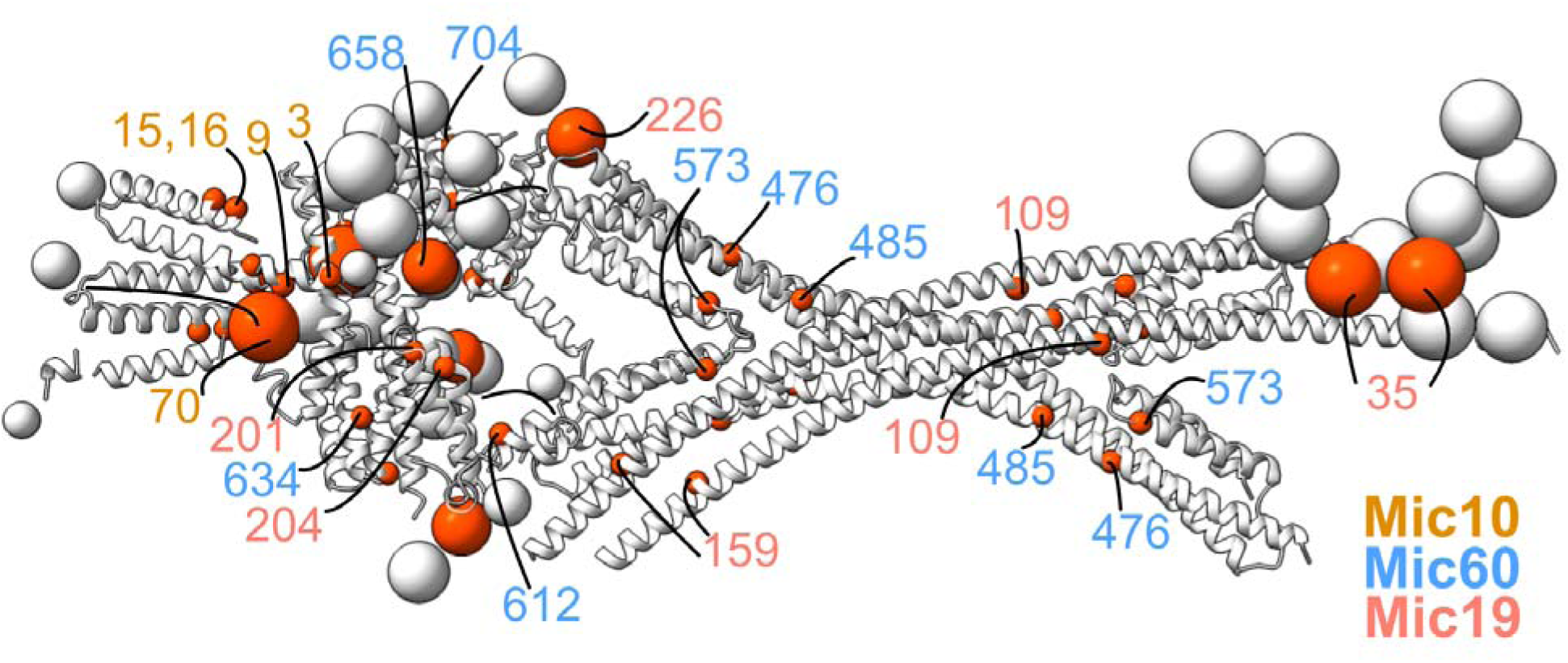
Mapping mutations from ClinVar on the MICOS integrative structure. ClinVar missense mutations of uncertain clinical significance, which are likely pathogenic based on AlphaMissense, are mapped to the representative cluster center bead model. The representative cluster center bead model is shown in grey, with the mutated residues colored in red. The residues are labeled in colors based on the protein. For a detailed list of mutations, see Table S8.

## Discussion

In this study, information from XLMS, biochemical assays, ET, AF predictions, homology modeling, and sequence alignments is used to obtain the integrative structure of the 4:2:2:1 Mic60^IMS^:Mic19:Mic10:Mic13 MICOS sub-assembly corresponding to the MICOS subunits in one half of the CJ, *i.e.,* a half-junction of MICOS, at 16 Å precision. As shown in the schematic, the tetrameric Mic60^CC^, which follows its N-term disordered region and Mic60^HB^, extends into the CJ, connected to the membrane-proximal domains (Mic60^LBS1,LBS2,mitofilin^) by a small helical region, Mic60^link^ (Fig. 7). Next, in Mic19, the Mic19^N^ extends above the CJ, allowing it to interact with the OMM complexes (Fig. 3D) [38], [39]. The disordered last 20 residues of Mic19^N^ wrap around the parallel Mic19^CC^ dimer. Together with Mic60^CC^, Mic19^CC^ is thought to form a selective barrier in the CJ, regulating the passage of proteins and metabolites [5], [22]. The Mic19^CHCH^ domain, which follows Mic60^CC^, binds to the membrane-proximal domains of Mic60 (Mic60^LBS1,LSB2,mitofilin^). Next, in Mic10, Mic10^N^ is mostly localized above the Mic60^mitofilin^. The two transmembrane helices of a Mic10 monomer adopt a wedge-shaped conformation in the integrative structure as previously hypothesized [15], [16]. The two monomers in a Mic10 dimer, oriented at an angle with respect to each other, bind *via* their respective Mic10^TM2^ helices. Together, these intra- and inter-Mic10 monomer angles may induce membrane curvature, providing the structural basis for membrane bending by Mic10 homo-oligomers. Mic10^C^, which follows Mic10^TM2^, interacts with the Mic13^central,C^, consistent with the *in vitro* experiments in this study, which demonstrate that Mic10^C^ binds to Mic13, whereas Mic10^N^ does not. Finally, in Mic13, Mic13^TM^ binds to Mic10^TM1^, and is followed by Mic13^central^, an amphipathic helix mostly parallel to the IMM, and Mic13^C^, which localizes below Mic60^mitofilin^ (Fig. 4A, Fig. 7).

**Figure 7.**
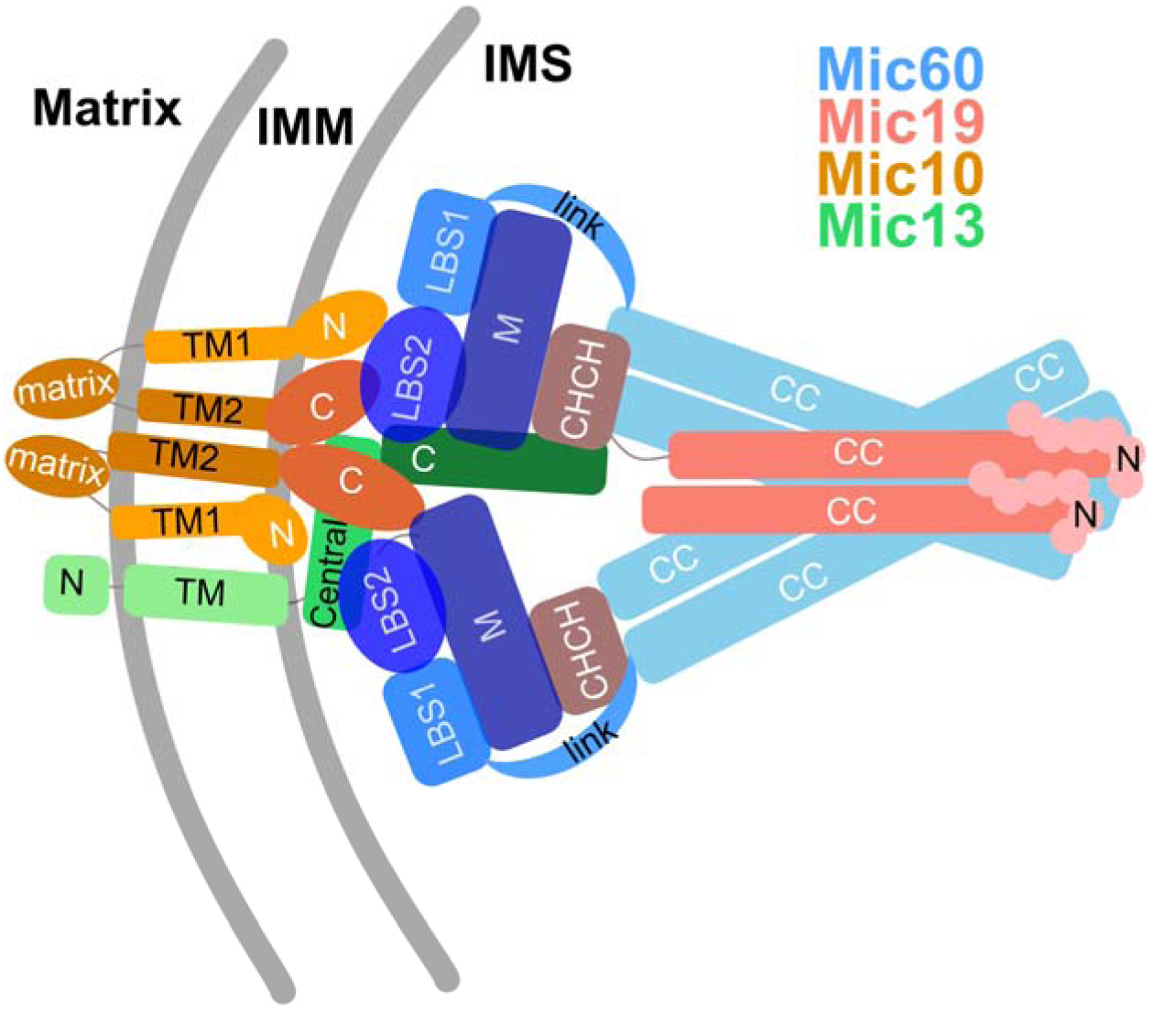
Proposed schematic of the MICOS sub-assembly. Based on the integrative structure, a proposed architecture of the MICOS sub-assembly, containing Mic10, Mic13, Mic19, and Mic60, is shown. The view from the top of the CJ is shown.

Mic13 has been postulated as a bridge from earlier studies [30], [40], [41], although recent studies have disputed it showing its role is to stabilize Mic10 [20]. Our integrative structure suggests that Mic13^central,C^ interacts with the Mic10^C^ and Mic60^LBS1,LBS2,mitofilin^ regions (Fig. 4). Further, *in vitro* experiments show direct interactions between Mic13-Mic10^C^ and Mic60^IMS^-Mic13 (Fig. 5). Together, these results indicate that Mic13 might play a role in connecting the Mic10- and Mic60-subcomplexes during MICOS assembly. Mic13 acts as a bridge by associating with Mic10 predominantly in the IMM, and with Mic60 in the IMS. We propose that the Mic13^central^ amphipathic helix acts as a flexible hinge that connects the Mic10-associated and Mic60-associated regions of Mic13. Although Mic13^central^ is heterogeneous in our integrative structure, it is unclear whether the heterogeneity is indeed due to underlying flexibility or a lack of data on this region. Consistent with our observation that Mic13 might play a role in bridging the Mic10- and Mic60-subcomplexes, deletion of Mic13 leads to reduced association between the two subcomplexes, altered cristae morphology, and a concomitant reduction in the number of cristae and cristae junctions [20], [30], [40], [41].

The structure of Mic19^CC^ was hitherto unknown. Mic19^59-174^ (Mic19^CC^) is predicted to be a coiled-coil, according to sequence-based predictions using COILS [46], PCOILS [47], and MARCOIL [48]. It is expected to form a dimer, based on a previous *C. thermophilum* study and the presence of intra-Mic19 crosslinks between identical residues in this region [17]. In the integrative structure, Mic19^CC^ forms a parallel dimer. However, ∼30% of intra-Mic19 crosslinks are violated in this parallel coiled-coil configuration (Table S6). We investigated the fit to intra-Mic19 crosslinks by modeling both parallel and anti-parallel configurations of the Mic19^CC^ dimer. Notably, the intra-Mic19 crosslink satisfaction is slightly worse in the anti-parallel configurations (Table S10). This suggests that Mic19^CC^ may assemble into higher-order oligomers or adopt a folded conformation, which remains to be explored.

Next, we discuss the limitations of our study. In the present study, the stoichiometry of a minimal MICOS sub-assembly was assumed, based on known X-ray structures and AlphaFold predictions [22]. The N-terminal region of Mic60, Mic60^1-450^, and other MICOS subunits, such as Mic25 (a paralog of Mic19), Mic26, and Mic27, were not modeled due to the lack of interaction data on these proteins. Further, the Mic60^1-45^ residues are also not conserved between humans and fungi (average ConSurf score of 3.5). SLP2 and components of the MIB-SAM complex were also not modeled, although they may influence the MICOS architecture [8], [20], [29], [49].

We also considered modeling the holo complex at the CJ using symmetry to place the membrane-proximal domains of the modeled sub-assembly on the diametrically opposite sides of the CJ. However, the resulting complex will still not be equivalent to the holo complex: the previously mentioned unmodeled regions, such as Mic60^1-450^, cannot currently be incorporated, as there is no data to restrain them relative to the modeled sub-assembly (Nathanail et al., 2026). Also, given that several regions in the current model are already at low precision due to lack of experimental data, *e.g.*, Mic13^C^ and Mic19^CC^, adding symmetric copies would introduce additional uncertainty and further reduce the model precision (Fig. 3, Fig. S7). Moreover, small proteins such as Mic10 and Mic13 are not expected to span the symmetric axis; hence, interfaces related to these proteins would remain the same in the symmetric holo complex. Therefore, instead of modeling the holo complex, we decided to focus our study on determining the interactions between subunits in the modeled sub-assembly.

Our study constitutes the first step towards determining the complete molecular architecture of cristae junctions, including the spatial organization of MICOS complexes along the junction, and their association with the larger MIB complex. In this study, we determined the integrative structure of a MICOS sub-assembly comprising Mic10, Mic13, Mic19, and Mic60, where we identified and validated several novel interfaces among the MICOS subunits. Further experiments, such as quantitative mass spectrometry, can reveal the stoichiometry of subunits in the MICOS complex; X-ray crystallography and single-particle cryo-electron microscopy (cryo-EM) can provide high-resolution structures of the MICOS subunits and sub-complexes. The integrative modeling framework used here is inherently adaptable, allowing the structure to be updated iteratively, based on new data, such as stoichiometry information and new experimental structures.

## Methods

Integrative structure determination of the MICOS sub-assembly was performed in four stages (Fig. 1-2) [24], [25]. Here, we combined information from AF predictions, crosslinking mass spectrometry (XLMS), biochemical assays, electron tomography (ET), homology modeling, and sequence alignments to obtain the integrative structure of the Mic60^IMS^:Mic19:Mic10:Mic13 MICOS sub-assembly. We used the Python Modeling Interface (PMI) of the Integrative Modeling Platform (IMP 2.23.0; https://integrativemodeling.org. The pipeline is based on previous studies [32], [34]. UCSF Chimera v1.8 and UCSF ChimeraX v1.8 were used for visualization [50], [51]. The input data, scripts, and results are available at https://github.com/isblab/micos and (10.5281/zenodo.18383758). The integrative structures will be deposited in the PDB.

### Stage 1. Gathering information

#### Stoichiometry

We modeled a 4:2:2:1 Mic60^IMS^:Mic19:Mic10:Mic13 MICOS sub-assembly. The stoichiometry of Mic60 and Mic19 was based on the available X-ray structures of a Mic60^451-597^ coiled-coil homo-tetramer, and a 2:2 Mic60^627-648,684-752^:Mic19^178-221^ hetero-tetramer [22]. Mic10 exists as an oligomer in the MICOS complex [1], [15], [16]. Given the lack of experimental structures of Mic10, we used AlphaFold3 (AF3) to predict the structures of Mic10 homo-oligomers. We included two copies of Mic10 and one copy of Mic13 in our integrative structure, based on a confident 2:1 Mic10^TM^: Mic13^ΤΜ^ AF3 prediction.

#### Membrane dimensions

The membrane thickness was 4 nm, and the CJ diameter was 25 nm from electron tomography [9], [52].

#### Membrane topology

Information from co-IP, pull down assays, protein subcellular fractionation, and liposome sedimentation assays was used to determine the membrane topology of each subunit domain. This was used to restrain the corresponding model regions to the IMS, IM, and matrix regions (Fig. 1-2, Table S1, Table S5) [15], [16], [29], [30], [38], [39], [40].

#### Atomic structures

##### Homology modeling

The structures of human Mic60^451-597^ (coiled-coil tetramer) and Mic60^627-648,684-752^:Mic19^178-221^ hetero-tetramer were modeled using MODELLER based on the corresponding templates from *L. thermotolerans* and *C. thermophilum* (PDB: 7PUZ, 7PV1), respectively (Fig. 1-2) [22], [53].

##### AlphaFold predictions

We used AF3 extensively for modeling several combinations of the MICOS subunits (Table S1-S2). Each AF3 prediction was performed with 20 seeds (100 models per prediction). The best-ranking model from each prediction was selected based on the ranking score. Predictions with an ipTM greater than 0.8 were considered to be confident and were considered for further analysis. Confident regions were extracted from these AF3 predictions using a graph-based community clustering method on the PAE matrix to define pseudo-domains (PAE cutoff of 12), followed by filtering the residues in these pseudo-domains with a pLDDT threshold of 70, as implemented in AF-pipeline [54], [55]. To increase the structural coverage, we included additional structures of individual subunit domains from the AlphaFold database, from which confident regions were extracted using the same criteria (Table S1-S2) [56].

##### Chemical crosslinks from mass spectrometry

Data from XLMS, obtained from the XLinkDB and several other studies, were used to restrain the distances between the crosslinked residues in the modeling, as well as for validating the model [57], [58], [59], [60], [61], [62], [63], [64]. We obtained a total of 341 crosslinks from human and mouse studies corresponding to a variety of crosslinkers: 68 BDP, 73 PIR, 6 DHSO, 113 DSSO, and 81 BS3 (Table S4). The crosslinks obtained from mouse studies were mapped to human subunits, and the mouse crosslinks on Mic25 were mapped to the homologous human Mic19, based on multiple sequence alignments (MSA) using MAFFT [65]. Of these, 303 crosslinks mapped to residue pairs in the MSAs and 38 crosslinks mapped to gaps. Around 80% (239) of the 303 crosslinks were used in sampling, and the remaining 20% were used for validation.

##### Protein-protein binding assays

Data on interacting protein domains from co-immunoprecipitation (co-IP), immunoblotting, blue native PAGE (BN-PAGE), and pull-down assays were used to restrain the corresponding distances between these domains in the model (Table S3) [15], [16], [28], [29], [30].

### Stage 2. Representing the system and translating the data into spatial restraints

#### Subunits and number of copies

We modeled the MICOS sub-assembly with 4, 2, 2, and 1 copies of Mic60^451-597^, Mic19^1-227^, Mic10^1-78^, and Mic13^1-118^, respectively.

#### Multi-scale coarse-grained bead representation

We used a multi-scale coarse-grained bead model to represent the MICOS subunits, with each protein represented as a string of spherical beads. Domains with known atomic structures were represented as a rigid body using a multi-scale representation at one and ten residues per bead to maximize computational efficiency. Domains with unknown structures were represented at a coarser resolution of 10 residues per bead (5 residues per bead for the Mic60^LBS2^ domain) and allowed to move as flexible strings of beads.

A subset of the gathered information was encoded into spatial restraints described below. These restraints together constitute a Bayesian scoring function that allows for sampling models that are consistent with the input information [66].

#### Bayesian crosslinking restraints

The distances between the crosslinked residues were restrained using the Bayesian crosslink restraint [66]. The restraint uses a compound likelihood term that considers multiple residue pairs assigned to a single crosslink to account for ambiguity (multiple copies of a subunit). It also accounts for the uncertainty in the position of the crosslinked residues and uncertainty in the crosslinked length due to backbone and side-chain fluctuations, apart from other uncertainties associated with crosslinks, such as the false positive rate of the crosslinking experiment and the ambiguity in the assignment of a crosslink due to multiple copies of subunits.

#### Protein-protein interaction restraints

Distances between interacting protein domains (from biochemical studies) and contact residue pairs (from AF3 predictions) were restrained using minimum pair restraints (Table S3, S9). This is a harmonic upper-bound restraint on the maximum distance between pairs of beads representing the two interacting protein domains in a model. To account for ambiguity or multiple copies of the same protein, distance restraints for two interacting proteins, A and B, were applied such that at least one copy of protein A was restrained to be close to at least one copy of protein B.

#### Membrane restraints

Based on the membrane dimensions, the crista was modeled as a cylinder centered at the origin, with its base parallel to the XY plane and its axis parallel to the Z axis. The inner and outer radii of the cylinder were set to 12.5 nm and 16.5 nm, respectively (Fig. S1, Table S5) [9], [52]. The following restraints were used to restrain the beads according to their membrane topology.

#### Transmembrane restraints

The beads corresponding to the transmembrane domains were restrained to be in the IMM region, *i.e.,* between the outer and inner radii of the cylinder. The restraint was implemented using harmonic upper and lower bounds on the radial distance of each bead from the cylinder axis.

#### IMS localization restraints

The beads corresponding to the IMS domains were restrained to be in the IMS region, *i.e.,* inside the inner radius of the cylinder. The restraint was implemented using a harmonic upper bound on the radial distance of each bead from the cylinder axis.

#### Matrix localization restraints

The beads corresponding to the matrix domains were restrained to be in the matrix region, *i.e.,* outside the outer radius of the cylinder. The restraint was implemented using a harmonic lower bound on the radial distance of each bead from the cylinder axis.

#### Z axial restraints

The beads corresponding to Mic10 and Mic13 were restrained to remain close to the upper rim of the crista cylinder, representing the crista junction. The beads corresponding to the Mic19^1-14^ residues were restrained to be above the crista junction, as these are known to interact with Sam50 in OMM [38], [39]. Finally, the beads corresponding to the Mic60^coiled-coil^ tetramer were restrained at the upper rim of the cylinder based on the dimensions of the tetramer and the CJ [22]. The restraint was implemented using a harmonic upper and lower bound only along the Z axis.

#### Stereochemistry restraints

The excluded volume restraint was used to avoid steric clashes by preventing beads from overlapping. The restraint was encoded as a soft-sphere pair score that penalizes the distances between beads if they are smaller than the sum of their radii [24]. The sequence connectivity restraint was used to restrain the distance between consecutive beads in a protein. The restraint was encoded as a harmonic upper bound score that penalizes the distances between beads that are greater than a threshold distance.

### Stage 3. Configurational sampling

We followed the Gibbs sampling Replica Exchange Markov Chain Monte Carlo algorithm for the positions and orientations of the rigid bodies and flexible beads [35], [37]. We started with initial random configurations for all domains, except for the Mic60-Mic19 tetramer. The latter is fixed at the IMM, positioned according to the lipid-binding sites in Mic60 [22]. The beads localized in the IMS, IMM, and matrix were shuffled in their respective bounding boxes, defined based on the membrane topology (Fig. S1). We performed 50 independent runs with 8 replicas and 50000 frames per run, sampling 200 million models.

### Stage 4. Analysis and validation

We followed the analysis and validation protocol as used in previous studies [32], [34], [35], [37], [67], [68]. First, the sampled models were analyzed to assess the sampling exhaustiveness, including clustering and estimation of the model precision. We obtained a single major cluster of models, which represents the outcome of integrative modeling, and is hereafter referred to as the integrative model.

Next, we assessed the fit of the models to the data used in the modeling, as well as to the data not used in modeling. Subsequently, we analyzed protein-protein interfaces in the model; a subset of them was also validated by *in vitro* experiments. Finally, we rationalized missense mutations using the integrative model. The integrative model was visualized in two ways: as a bead model representing the cluster, and as a localization probability density (LPD) map. The LPD map shows the probability of a voxel being occupied by bead(s) in superposed models. We also used PrISM to annotate high- and low-precision regions in the integrative model [36].

#### Fit to input data

First, we calculated the fit to the input interacting protein domains (from biochemical studies) and contact residue pairs (from AF3 predictions) by computing the minimum distance across all the interacting pairs of beads, for each model in the cluster (Fig. 2, Fig. S4).

Next, we calculated the fit to the input crosslink data by computing the distance between crosslinked residues in each model in the cluster. A crosslink is said to be satisfied if the minimum crosslink distance in at least one model in the cluster is less than the cutoff, set to 52 Å (BDP/PIR), 35 Å (BS3), and 30 Å (DHSO/DSSO) (Fig. 2, Fig. S5, Table S6).

Lastly, we calculated the fit to the membrane localization data by computing the corresponding restraint violations for models in the cluster.

#### Fit to data not used in the modeling

The fit to crosslinks not used in the modeling was calculated in the same way as fit to the input crosslinks (Fig. 2, Fig. S6, Table S6).

#### Contact maps

To identify the protein-protein interfaces, we computed regions of contact between each protein copy pair. These regions comprise bead pairs in contact; a bead pair is in contact if the average distance between their surfaces across models in the cluster is less than or equal to 10 Å.

#### Mapping missense variants

We obtained missense mutations labeled pathogenic or of uncertain significance on the modeled MICOS subunits from ClinVar [43]. Subsequently, we used AlphaMissense to assign a pathogenicity score to these mutations [42]. The mutations annotated as pathogenic (AlphaMissense score >0.5) were mapped on the integrative structure.

### Recombinant Protein Purification and Fluorophore labelling

The gene encoding wild type ctMic12 (ctMic12^WT^) was cloned into a pET28a(+) vector as an N-terminal SUMO fusion construct and transformed into *Escherichia coli* Rosetta (DE3) cells. Briefly, *E. coli* Rosetta (DE3) cells expressing ctMic12 were grown in terrific broth at 37 °C to an OD□□□ of ∼0.8 and induced with 0.1 mM IPTG, followed by overnight expression at 18 °C. Cells were lysed by sonication in buffer containing 50 mM Tris-HCl (pH 8.5), 500 mM NaCl, 10% (v/v) glycerol, 20 mM imidazole, and either 1% (w/v) n-dodecyl-β-D-maltoside (DDM) or 2% (w/v) sarkosyl. The lysate was clarified and applied to TALON (Co²□) affinity resin. The resin was washed with a wash buffer containing 50 mM Tris-HCl (pH 8.5), 500 mM NaCl, 10% (v/v) glycerol, 20 mM imidazole, and 0.05% DDM. Bound protein was eluted in a buffer containing 10 mM Tris-HCl (pH 8.5), 150 mM NaCl, 5% (v/v) glycerol, 0.05% DDM, and 350 mM imidazole. The SUMO tag was cleaved by the Ulp1 protease (1:100, w/w) at 4 °C overnight. The cleaved protein was further purified by size-exclusion chromatography (SEC) in buffer containing 10 mM Tris-HCl (pH 8.5), 150 mM NaCl, 5% (v/v) glycerol, and 0.05% DDM, concentrated to ∼1 mg/mL, flash-frozen in liquid nitrogen, and stored at −80 °C until further use.

To enable site-specific fluorophore labelling, a S62C variant of ctMic12 (ctMic12^S62C^) was generated by site-directed mutagenesis to introduce a cysteine residue. For fluorophore labelling, purified ctMic12 S62C was buffer exchanged into 20 mM HEPES (pH 7.0), 150 mM NaCl, 5% (v/v) glycerol, 0.05% DDM, and 0.5 mM TCEP, and incubated with Alexa Fluor 488 C5-maleimide at a 1:20 molar ratio (protein:dye) for 3 h at room temperature. Excess dye was removed, and buffer exchange was performed in a single SEC step into buffer containing 10 mM Tris-HCl (pH 8.5), 150 mM NaCl, 5% (v/v) glycerol, and 1% DDM. The concentration of Alexa Fluor 488-conjugated ctMic12 S62C was estimated using a fluorescence-based plate reader assay with Alexa Fluor 488 as the standard, and 300 μL aliquots were snap-frozen and stored at −80 °C.

A previously reported construct of ctMic60^IMS^ (residues 208-691) was expressed and purified as described [22]. Briefly, *E. coli* BL21 (DE3) cells expressing ctMic60 were grown in terrific broth at 37 °C to an OD□□□ of ∼0.8 and induced with 0.2 mM IPTG, followed by overnight expression at 18 °C. Cells were lysed by sonication in buffer containing 50 mM Tris-HCl (pH 8.0), 500 mM NaCl, 10% (v/v) glycerol, and 20 mM imidazole. The lysate was clarified and applied to a 5 mL HisTrap FF column (Cytiva) using an ÄKTA Pure system. After washing with lysis buffer, bound protein was eluted with a buffer containing 300 mM imidazole. The eluted protein was further purified by SEC, and the main peak fractions were pooled, concentrated to ∼5 mg/mL, flash-frozen, and stored at −80 °C. To enable downstream fluorophore labelling, a S2C mutant of ctMic60^IMS^ (ctMic60^IMS-S2C^) mutant was generated by site-directed mutagenesis and purified using the same protocol as ctMic60^IMS^. The purified protein was conjugated with Alexa Fluor 488 C5-maleimide using the same labelling conditions described for ctMic12^S62C^, except that DDM was omitted from the buffers. Excess dye was removed by size-exclusion chromatography.

### Synthetic Peptides

Peptides corresponding to the ctMic10^N^ and ctMic10^C^ regions of ctMic10 were designed with a 1× FLAG tag fused to either the N- or C-terminus, as appropriate, and commercially synthesized (sequences listed in Table S11). Lyophilized peptides were dissolved in Milli-Q water to a stock concentration of 1 mM and diluted to a 100 µM working concentration in MST buffer containing 10 mM Tris-HCl (pH 8.5), 150 mM NaCl, 5% (v/v) glycerol, and 0.05% (w/v) DDM.

### Microscale thermophoresis (MST) assays

Microscale thermophoresis (MST) experiments were performed using a Monolith NT.115 instrument (NanoTemper Technologies). For ctMic12-ctMic10 peptide and ctMic60^IMS^-ctMic10 peptide interaction studies, 25 nM Alexa Fluor 488-labelled ctMic12 and 5 nM Alexa Fluor 488-labelled ctMic60 were mixed with increasing concentrations of ctMic10^N^ or ctMic10^C^ in a total volume of 10 µL. The samples were then loaded into standard coated capillaries (NanoTemper Technologies). Measurements were carried out in triplicate at 25 °C using 20% and 70% excitation power and medium MST power settings. For ctMic12-Mic60^IMS^ protein-protein interaction studies, 5 nM Alexa Fluor 488-labelled ctMic12 was mixed with increasing concentration of Mic60^IMS^ in a total volume of 10 uL. Measurements were carried out in triplicate at 25 °C using 70% excitation power and medium MST power settings. Data analysis was performed using the NanoTemper analysis software.

### Circular dichroism (CD) measurements

CD spectra of 2-5 µM ctMic12 WT and the S62C variant were recorded using a JASCO spectropolarimeter with a 1 mm path-length quartz cuvette over the wavelength range of 200–260 nm. Three technical replicates were recorded for each sample and the baseline corrected CD spectra were plotted using Origin software.

## Data availability

Files containing input data, scripts, and results are available at https://github.com/isblab/micos and <u>10.5281/zenodo.18383758</u>. Integrative structures will be deposited in the PDB.

## Acknowledgements

We would like to thank Dr. Vinothkumar Kutti Ragunath for his input on AlphaFold predictions for membrane proteins, and Dr. Kiran Kulkarni and Dr. Pankaj Suman for discussions on the MST assay. We also thank ISB lab members Mubashira K.P. and Omkar Golatkar for helpful comments on the manuscript. The use of NCBS institutional computational and storage facilities is gratefully acknowledged. Molecular graphics images were produced using the UCSF Chimera and UCSF ChimeraX packages from the Resource for Bio-computing, Visualization, and Informatics at the University of California, San Francisco, supported by NIH R01-GM129325.

## Funding

S.V. acknowledges support from the Department of Atomic Energy (DAE), Government of India, under Project Identification No. RTI 4006 and RTI 4018, and from the Department of Biotechnology (DBT), Government of India, grant BT/PR40323/BTIS/137/78/2023. ATV acknowledges support from the Department of Atomic Energy (DAE), Government of India, under Project Identification No. RTI 4007, the Department of Biotechnology (DBT), Government of India for the Ramalingaswami Re-entry Fellowship (BT/RLF/Re-entry/69/2020) and the Science and Engineering Research Board (SERB/ANRF), Department of Science & Technology, Government of India via the Start-up Research Grant (SRG/2020/002578).

## Author contributions

**Muskaan Jindal:** conceptualization; methodology (computation); investigation; data curation; validation; visualization; writing─original draft; writing─review and editing; software.

**Rakesh Mahato:** conceptualization; methodology (experiments); investigation; validation; writing─original draft.

**Sreemoyee Das:** methodology (experiments); investigation; validation.

**Arko Guha:** methodology (experiments); investigation; validation; writing─original draft.

**Kartik Majila:** methodology (computation); software.

**Shreyas Arvindekar:** methodology (computation); software.

**Anand T. Vaidya:** conceptualization; methodology; investigation; validation; visualization; writing─original draft; writing─review and editing; funding acquisition; project administration; supervision; resources.

**Shruthi Viswanath:** conceptualization; methodology; investigation; validation; visualization; writing─original draft; writing─review and editing; software; funding acquisition; project administration; supervision; resources.

## Declaration of interests

The authors declare no competing interests.

## Declaration of generative AI and AI-assisted technologies in the writing process

During the preparation of this work, the authors used ChatGPT 5.6 (May-Oct 2026) to refine the text from initial drafts. After using these tools, the authors reviewed and edited the content as needed and take full responsibility for the content of the publication.

## Notes

### Competing Interest Statement

The authors have declared no competing interest.

### Summary of Updates

Updated to address reviewer comments from journal

