## Supplementary Material for "Integrative structure determination of a human mitochondrial contact site and cristae organizing system (MICOS) sub-assembly"

### Supplementary Figures

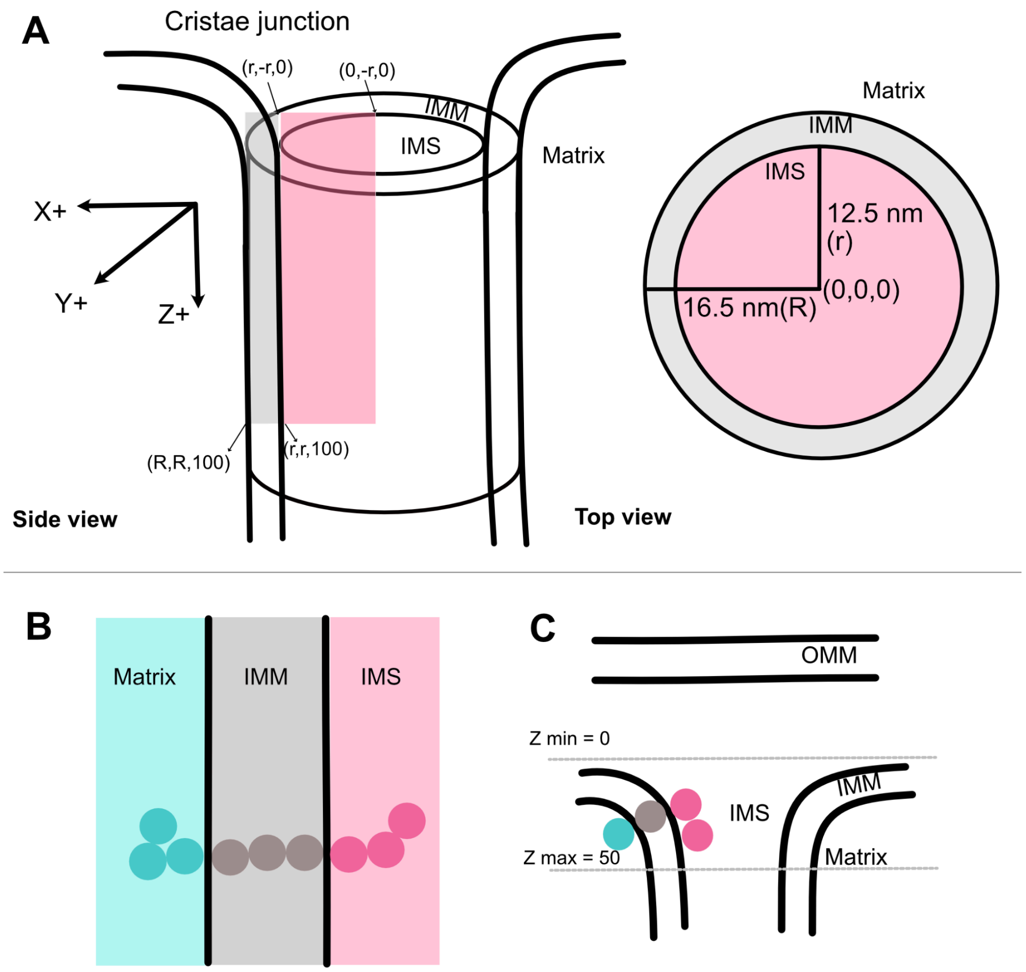


**Supplementary Figure 1. Membrane restraints used in the integrative modeling of the MICOS sub-complex. (A)** The cristae are modeled as hollow cylinders with inner (r) and outer (R) radii. Pink (grey) boxes show the bounding boxes for IMS (IMM) regions. **(B)** Membrane restraints are applied to restrict IMS-domain beads to the IMS region, TM-domain beads to the IMM region, and matrix-domain beads to the matrix region, respectively. The membrane restraint is implemented by computing the radial distance $(rad)$ in the XY plane from the center $(0, 0, 0)$, $rad = \sqrt{x^2 + y^2}$, for a bead with coordinates $(x, y, z)$. **(C)** Z-axial restraints are applied to all the beads corresponding to the MICOS subunits to restrain them at the rim of the CJ. See also Table S4.

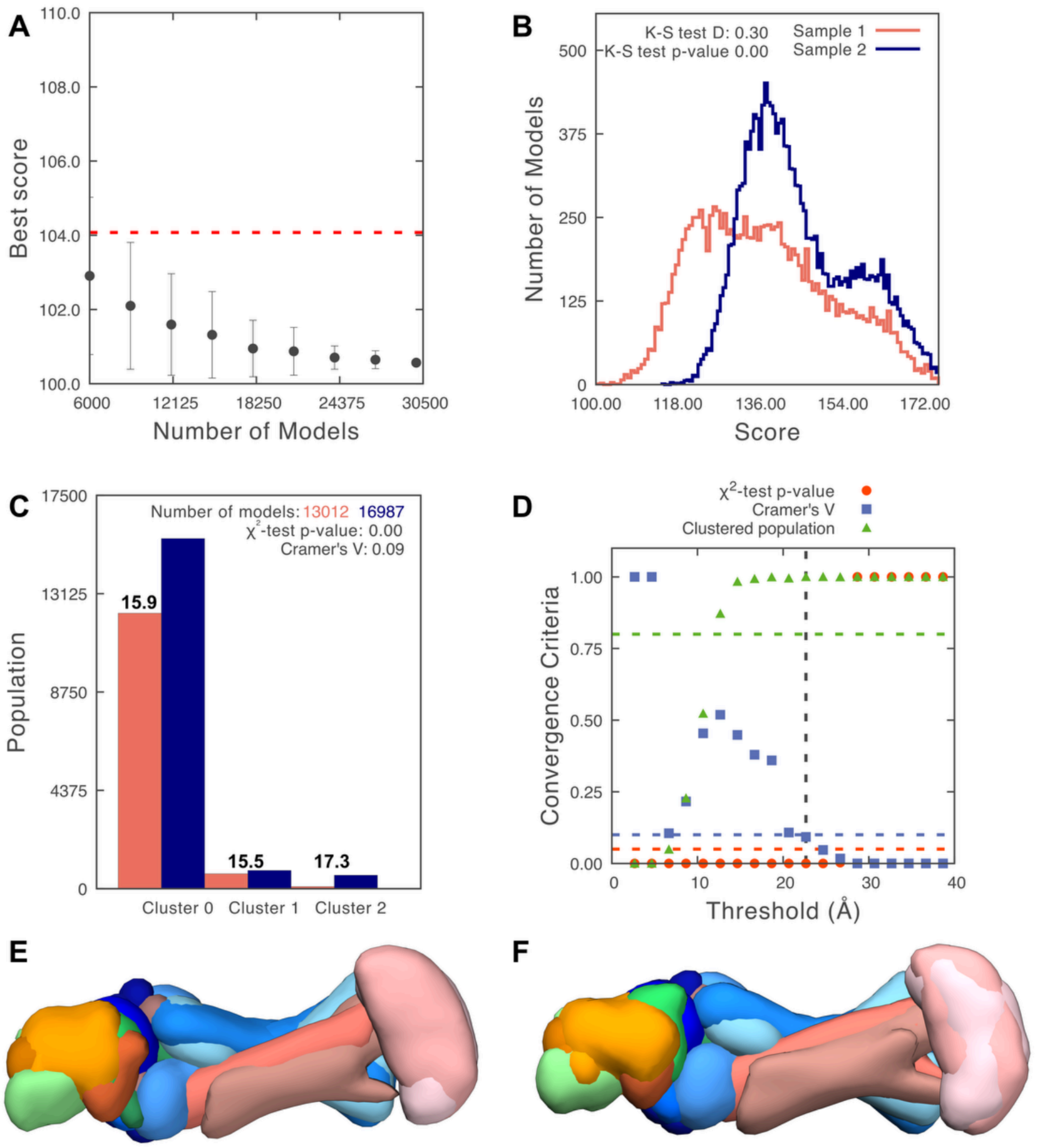


**Supplementary Figure 2. Sampling exhaustiveness protocol on MICOS subcomplex models. (A)** First test of convergence of the model score for the 28,773 good-scoring models. The scores are not improving further as more models are computed independently. The standard deviations of the best scores are plotted as error bars, which are estimated by repeating the sampling of models 10 times. The red dotted line depicts a lower bound reference on the total score. **(B)** Second test of similarity between the distribution of model scores of samples 1 (red) and 2 (blue). The difference in the distribution of scores is significant (Kolmogorov-Smirnov two-sample test p-value less than 0.05), but the magnitude of the difference is small (the Kolmogorov-Smirnov two-sample test statistic D is 0.3); thus, the two score distributions are effectively equal. **(C)** Populations of sample 1 and 2 models in the clusters obtained by threshold-based clustering using the RMSD threshold of 22.6 Å. Cluster precision is shown for each cluster. **(D)** Three criteria for determining the sampling precision (Y-axis), evaluated as a function of the RMSD clustering threshold (X-axis). First, the p-value is computed using the $\chi^2$-test for homogeneity of proportions (red dots). Second, an effect size for the $\chi^2$-test is quantified by the Cramer's V value (blue squares). Third, the population of models in sufficiently large clusters (containing at least 10 models from each sample) is shown as green triangles. The vertical dotted grey line indicates the RMSD clustering threshold at which three conditions are satisfied (p-value > 0.05 [dotted red line], Cramer's V < 0.10 [dotted blue line], and the population of clustered models > 0.80 [dotted green line]), thus defining the sampling precision of 22.6 Å. **(E-F)** The localization probability densities of models from sample A and sample B for the major cluster (97% population) were compared, and the cross-correlation between them is 0.97.

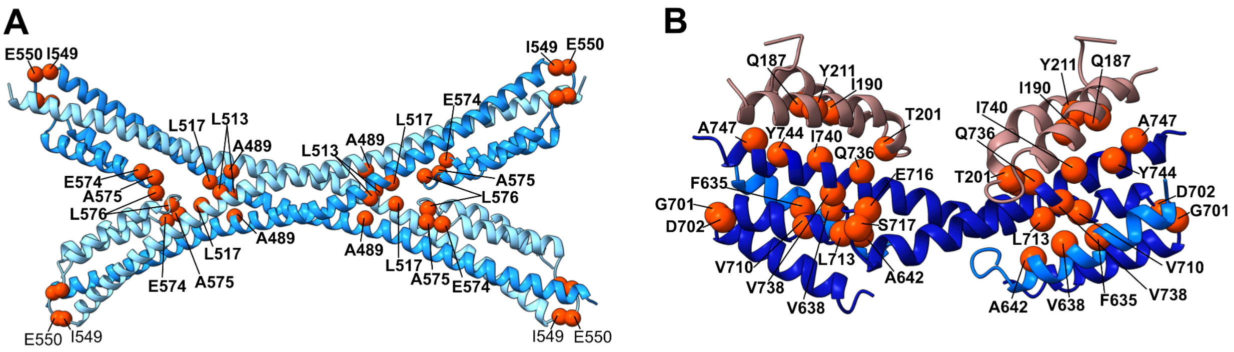


**C**

| Domain (residue-range) | Average ConSurf conservation score | Percentage of residues with >70% sequence conservation | Similarity percentage |
|---|---|---|---|
| Mic60$^{CC}$ (451-597) | 6.4 | 31 | 32 |
| Mic60$^{LBS1}$ (627-648) | 7.2 | 59 | 45 |
| Mic60$^{mitofilin}$ (684-751) | 7.1 | 54 | 56 |
| Mic19$^{CHCH}$ (178-221) | 6.3 | 54 | 40 |

**Supplementary Figure 3. Conserved residues mapped to the structures.** We used the multiple sequence alignment from a previous study [1], and the conserved residues in the interfaces and helix-break regions in **(A)** the Mic60$^{CC}$ and **(B)** Mic60$^{LBS1,mitofilin}$-Mic19$^{CHCH}$ structures are annotated as red spheres. **(C)** The modeled regions have ConSurf scores>6 [2] in the multiple sequence alignment across 30 species and >30% residues with >70% sequence conservation [1]. We also report the percentage of sequence similarity between the modeled human sequences and the corresponding sequences of the experimental structures, *i.e.,* Mic60$^{CC}$ from *L. thermotolerans*, Mic60$^{LBS1,mitofilin}$ and Mic19$^{CHCH}$ from *C. thermophilum*.

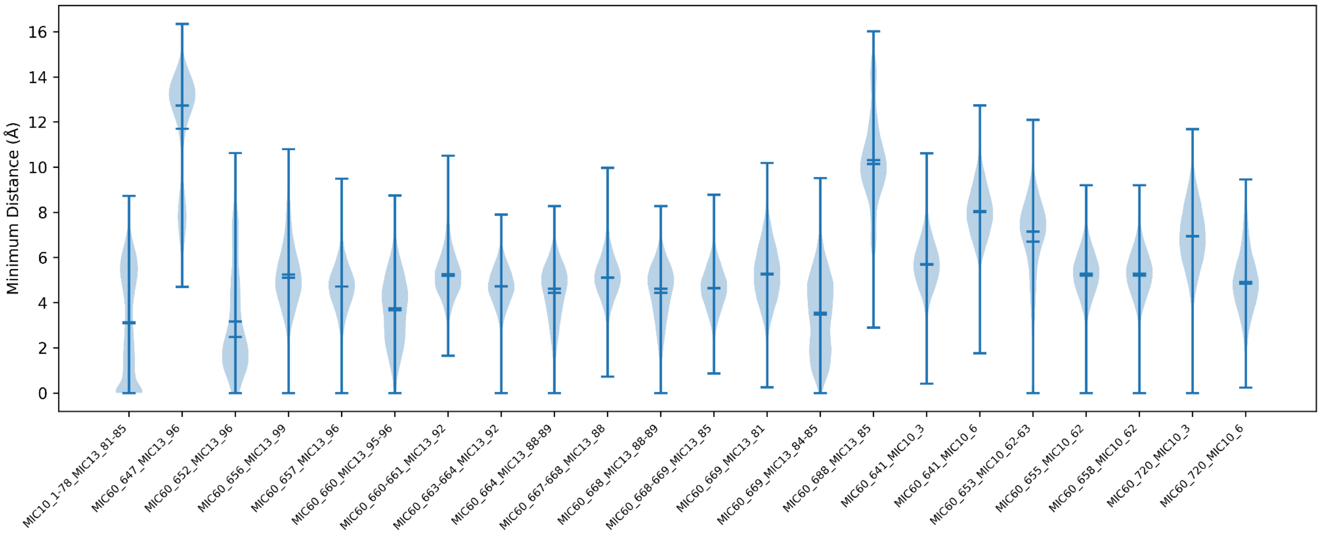


**Supplementary Figure 4. Fit to the biochemical data and AF3 interface residue predictions used in the modeling.** Violin plots showing the distribution of minimum distances between interacting protein domain pairs from biochemical data and AF3 interface residue predictions used in the modeling. The X-axis labels depict the interacting protein domains or residues in the form protein1_residue-range_protein2_residue-range.

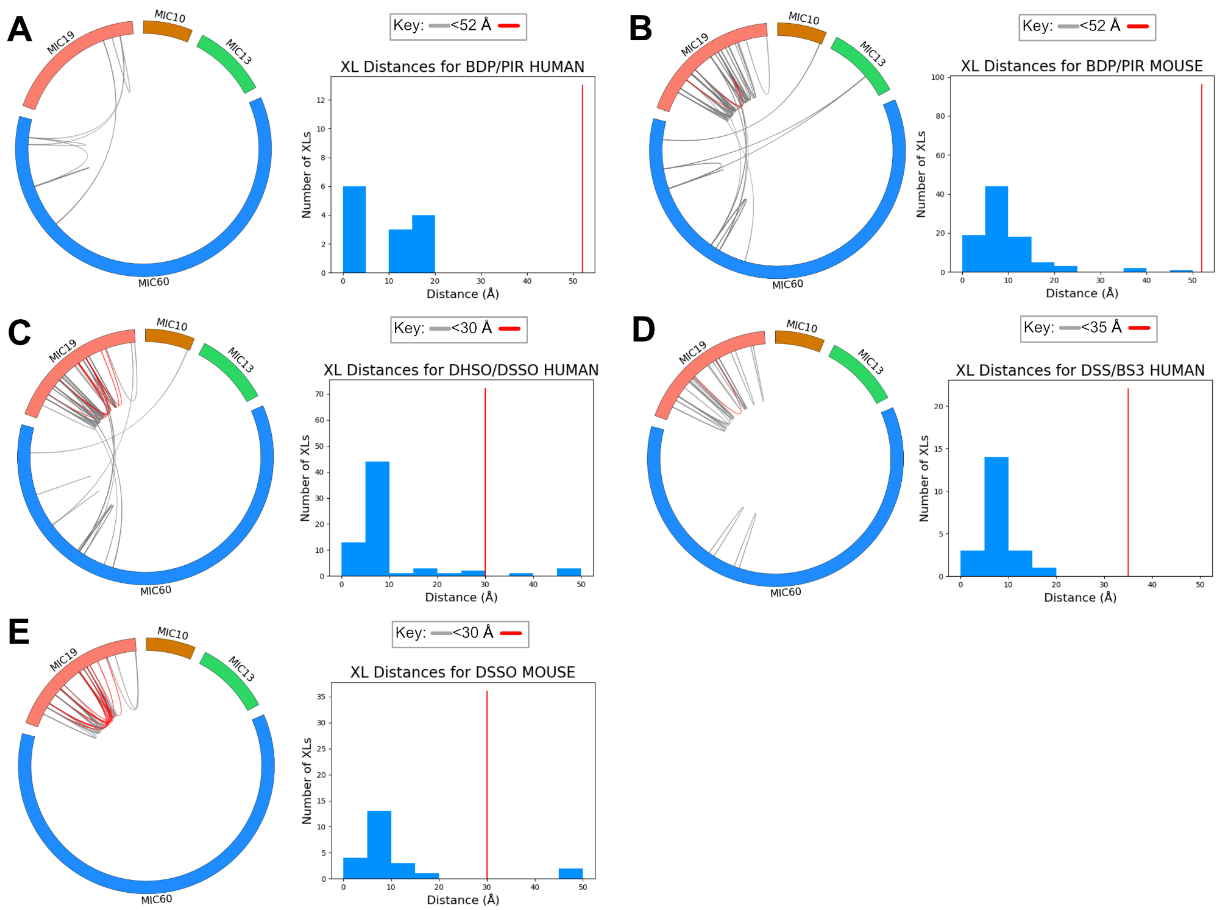


**Supplementary Figure 5. Results for the fit to XLMS data used in the modeling.** Circos plots from xiVIEW [3] and histograms show the fit to the XLMS data used in the modeling for **(A)** BDP/PIR human, **(B)** BDP/PIR mouse, **(C)** DHSO/DSSO human, **(D)** DSS/BS3 human, and **(E)** DSSO mouse crosslinks. A crosslink is considered satisfied by the cluster of integrative models if the minimum crosslink distance across all models in the cluster is within the violation threshold for the crosslinker type. The crosslinks in the Circos plots are colored grey (red) if they were satisfied (or not); the key mentions the threshold for a crosslinker. The histogram depicts these minimum crosslink distances, with the red line indicating the violation threshold. See also Table S7.

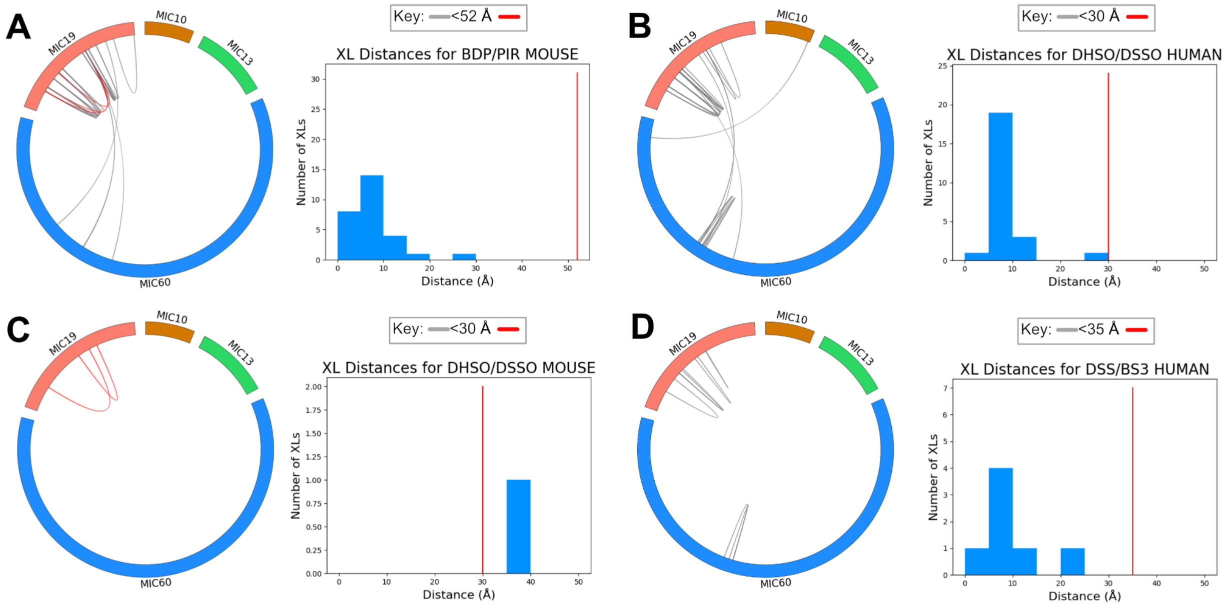


**Supplementary Figure 6. Results for fit to XLMS data not used in the modeling.** Circos plots from xiVIEW [3] and histograms showing the fit to the XLMS data used in the modeling, for **(A)** BDP/PIR mouse, **(B)** DHSO/DSSO human, **(C)** DHSO/DSSO mouse, and **(D)** DSS/BS3 human crosslinks. A crosslink is considered satisfied by the cluster of integrative models if the minimum crosslink distance across all models in the cluster is within the violation threshold for the crosslinker type. The crosslinks in the Circos plots are colored grey (red) if they were satisfied (or not); the key mentions the threshold for a crosslinker. The histogram depicts these minimum crosslinks distances, with the red line indicating the violation threshold. See also Table S7.

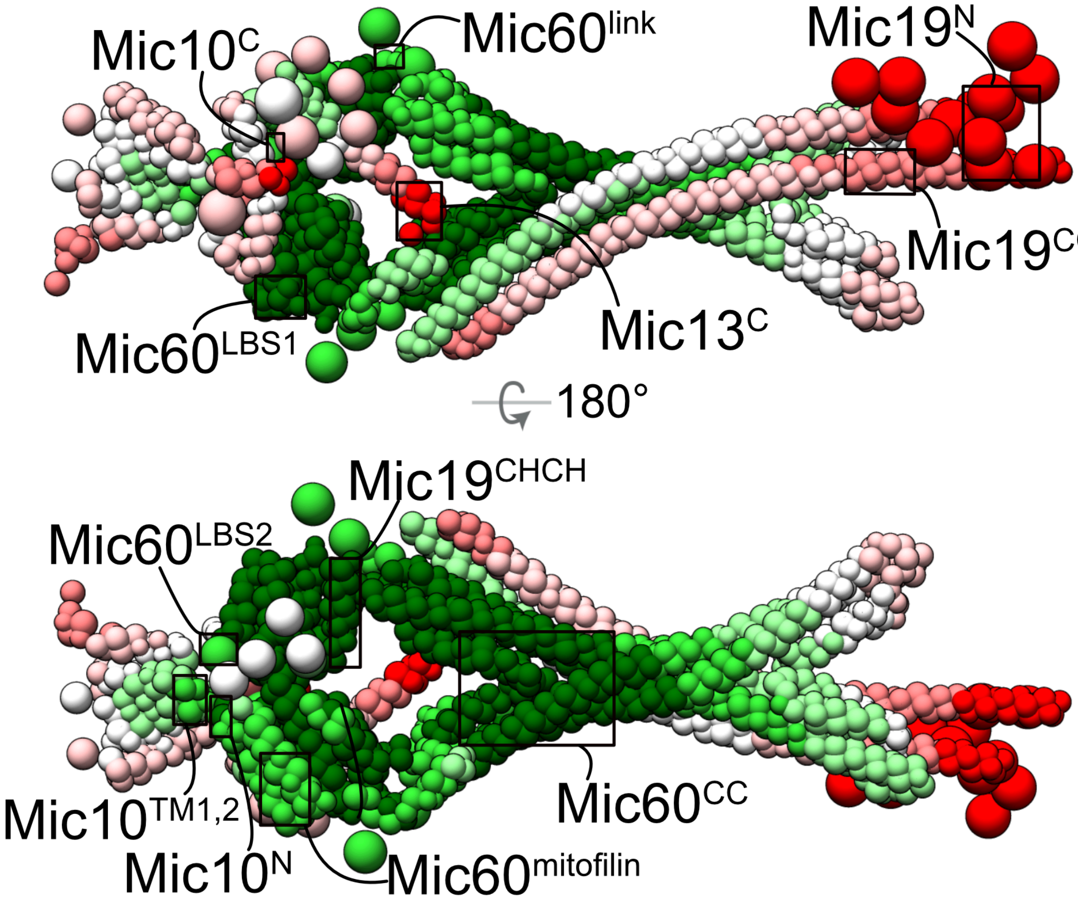


**Supplementary Figure 7. Integrative models annotated by PrISM.** The regions with high (green) and low (red) precision are marked on the representative bead model. See the Results section, "Integrative structure of a sub-complex of the MICOS complex."

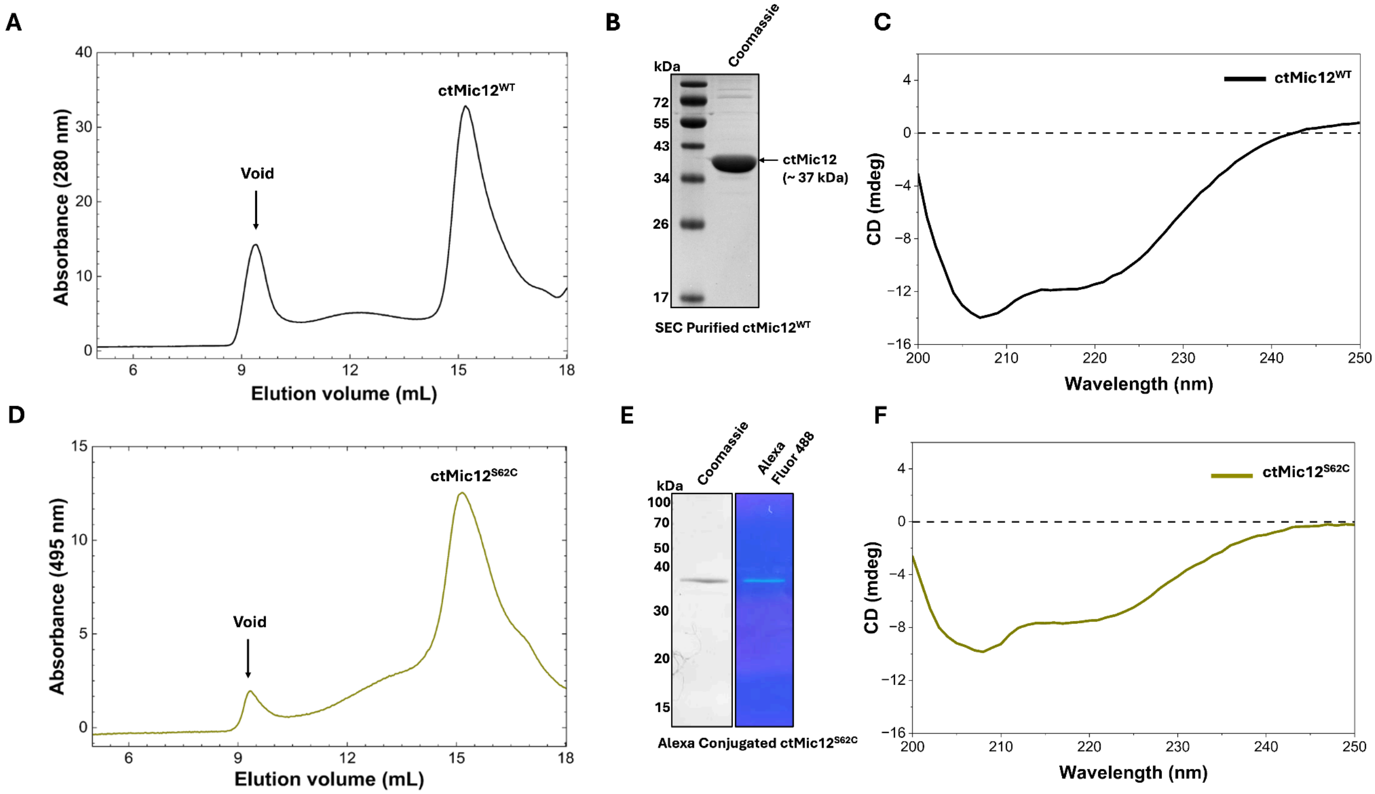


**Supplementary Figure 8. Purification and characterization of ctMic12 WT and the S62C variant. (A)** Size-exclusion chromatography (SEC) profile of ctMic12$^{WT}$. The peak fractions were used for further experiments. **(B)** Coomassie-stained SDS-PAGE gel showing purified ctMic12$^{WT}$. **(C)** Circular dichroism (CD) spectrum indicating predominantly α-helical secondary structure of ctMic12$^{WT}$. mdeg, millidegree of molar ellipticity **(D)** SEC profile of Alexa Fluor 488 C5-maleimide conjugated ctMic12$^{S62C}$, showing similar elution profile to ctMic12$^{WT}$ (A), suggesting that S62C substitution and fluorescent labelling do not alter its chromatography behaviour. The peak fractions were used for further experiments. **(E)** Coomassie staining (left) and UV illumination (right) of the SDS-PAGE gel showing Alexa Fluor 488-conjugated ctMic12$^{S62C}$. **(F)** CD spectrum of Alexa-conjugated ctMic12$^{S62C}$ indicating the preservation of its α-helical secondary structure.

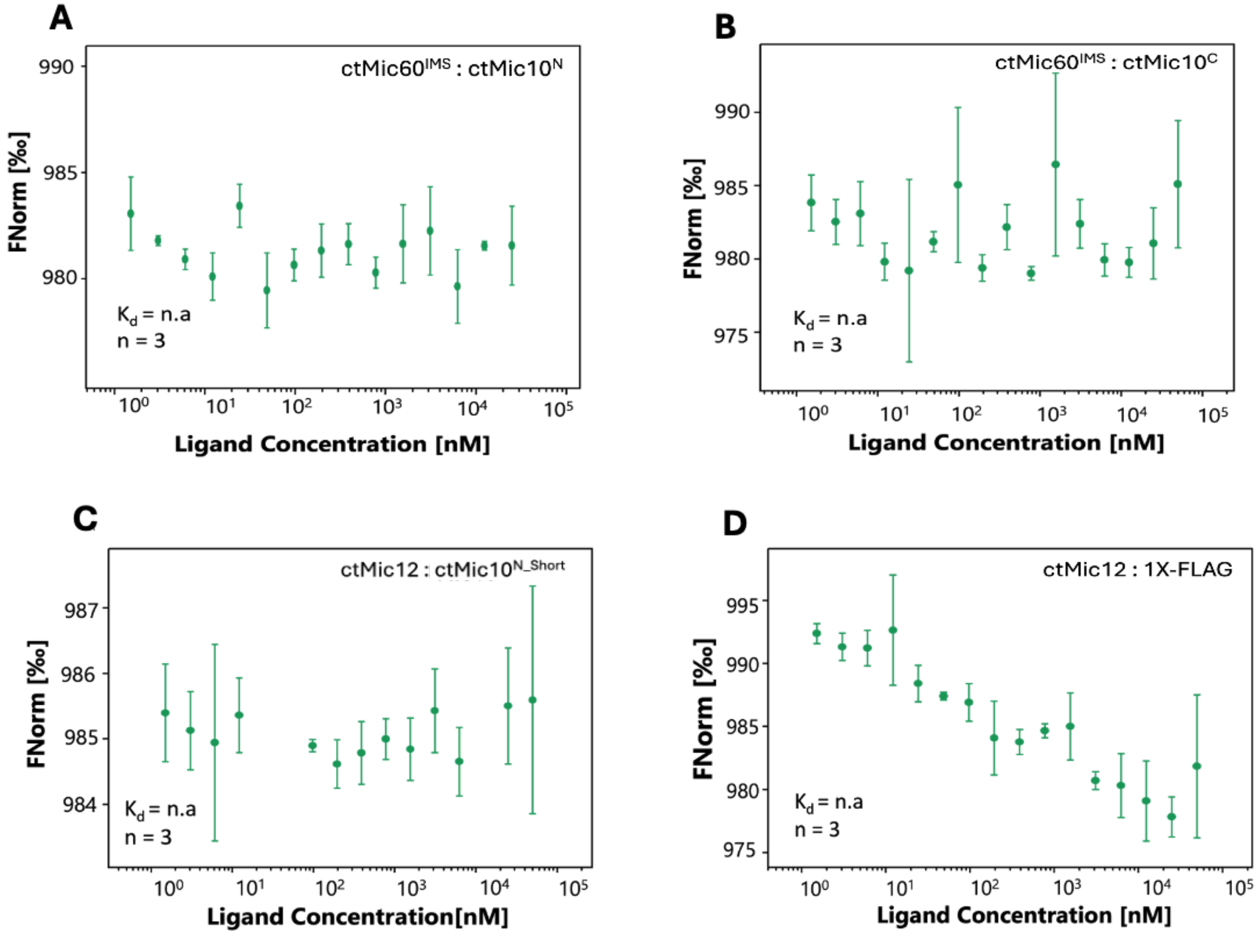


**Supplementary Figure 9. MST analysis of control interactions and Mic60 IMS binding with Mic10 peptides. (A–B)** MST analysis of ctMic60 IMS with the ctMic10$^{N}$ and ctMic10$^{C}$ showing no detectable interaction. **(C)** MST measurements of ctMic12 with a ctMic10$^{N_short}$ control peptide (sequence listed in Table S11), showing no detectable interaction. **(D)** MST measurements of ctMic12 with a 1× FLAG peptide control, indicating no detectable binding. In these measurements, ctMic60 S2C and ctMic12 S62C were fluorescently labeled and maintained at a constant concentration, while the partner protein/peptide was the ligand that was titrated. Normalized fluorescence (FNorm) is plotted as a function of ligand concentration. Error bars represent standard deviation from triplicate measurements.

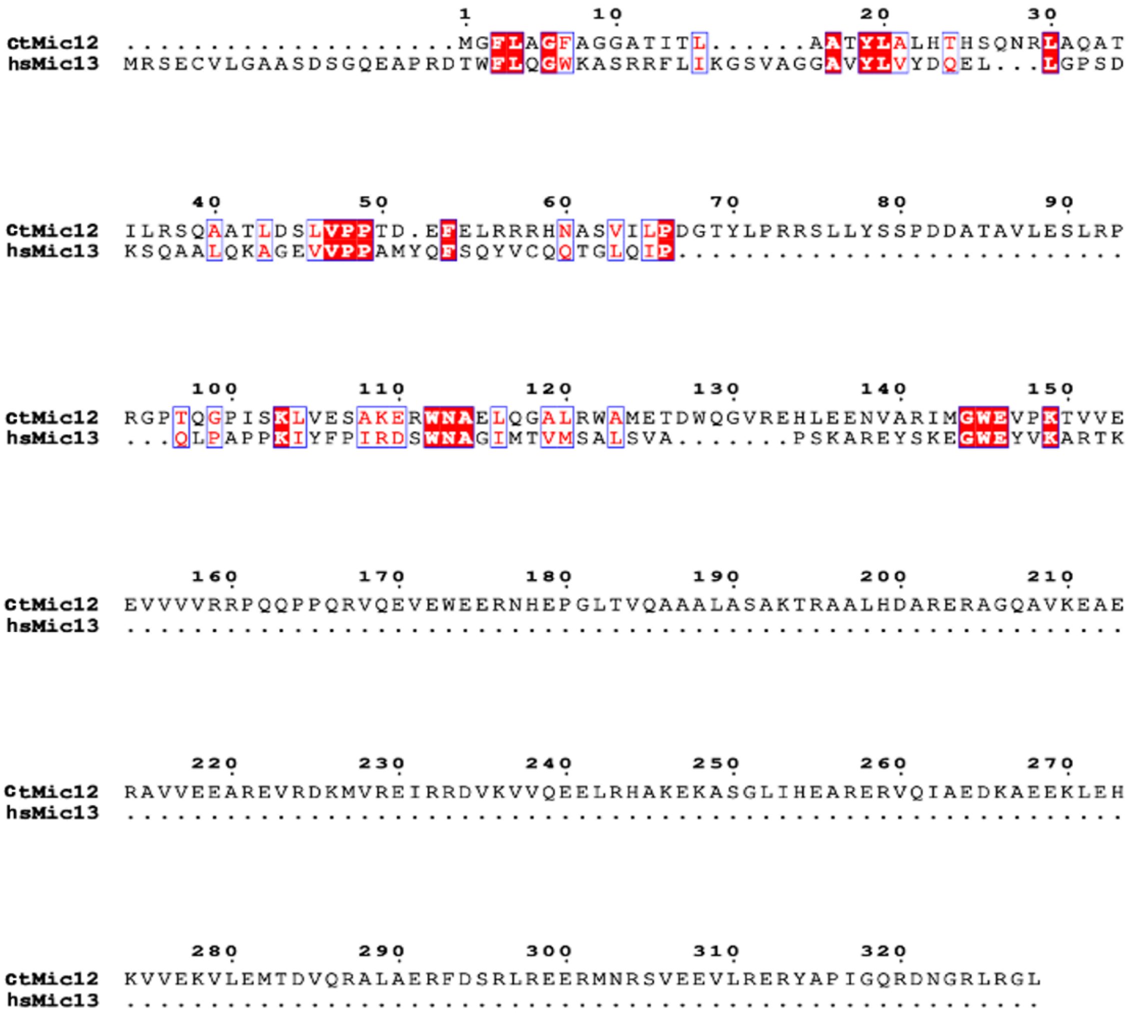


**Supplementary Figure 10. Sequence alignment of ctMic12 and HsMic13.** Sequence alignment of ctMic12 from *Chaetomium thermophilum* (ctMic12) and Mic13 from *Homo sapiens* hsMic13). Conserved residues are highlighted in red, with identical residues shown in white on a red background. Regions of sequence similarity are indicated, and residue numbering is shown above the alignment.

### Supplementary Tables

**Table S1. Structural information and membrane topology of the modeled subunits of the MICOS complex.** The domains for each protein are defined based on the membrane topology. The structure for each domain is obtained from AlphaFold (AF database or AF3 confident structure) or PDB. The Intermembrane space (IMS), Transmembrane (TM), and Matrix domains are highlighted in Pink, grey, and cyan, respectively.

| Protein | Uniprot ID | Residues | Structure | Domain name | Membrane topology |
|---|---|---|---|---|---|
| Mic10 | Q5TGZ0 | 2-12 | AF database | N | IMS |
| | | 13-36 | Mic10-Mic13 AF3 | TM1 | TM |
| | | 37-39 | Unknown | Matrix | Matrix |
| | | 40-60 | Mic10-Mic13 AF3 | TM2 | TM |
| | | 61-78 | Unknown | C | IMS |
| Mic13 | Q5XKP0 | 1-7 | AF database | N | Matrix |
| | | 8-23 | Mic10-Mic13 AF3 | TM | TM |
| | | 24-68 | AF database | Central | IMS |
| | | 69-78 | Unknown | Central | IMS |
| | | 79-118 | AF database | C | IMS |
| Mic19 | Q9NX63 | 1-58 | Unknown | N | IMS |
| | | 59-174 | AF database | Coiled-coil | IMS |
| | | 175-177 | Unknown | Central | IMS |
| | | 178-221 | Homology (PDB 7PV1) | CHCH | IMS |
| | | 222-227 | Unknown | CHCH | IMS |
| Mic60 | Q16891 | 451-597 | Homology (PDB 7PUZ) | Coiled-coil | IMS |
| | | 598-626 | AF database | Link | IMS |
| | | 627-648 | Homology (PDB 7PV1) | LBS1 | IMS |
| | | 649-683 | Unknown | LBS2 | IMS |
| | | 684-751 | Homology (PDB 7PV1) | Mitofilin | IMS |
| | | 752-758 | Unknown | Mitofilin | IMS |

**Table S2: AF3 predictions.** Monomer structures of the modelled MICOS subunits were obtained from the AlphaFold database. We obtained the confident regions from the AF predictions as described in the methods. PLM, OLA, and MYR refer to palmitic acid, Oleic acid, and Myristic acid. The rows highlighted in bold are used in integrative modeling.

| Protein copies | Lipids | Modeled domains | Modeled residue ranges | ipTM of the best ranking model |
|---|---|---|---|---|
| Higher-order oligomers | | | | |
| 4:2:2:1, Mic60-Mic19-Mic10-Mic13 | 10 PLM, 8 OLA, 8 MYR | Mic60$^{IMS}$, Mic19, Mic10, Mic13 | Mic60: 451-758, Mic19: 1-227, Mic10: 1-78, Mic13: 1-118 | 0.4 |
| 1:1:1:1, Mic60-Mic19-Mic10-Mic13 | 4 PLM, 4 OLA, 4 MYR | Mic60$^{IMS}$, Mic19, Mic10, Mic13 | Mic60: 451-758, Mic19: 1-227, Mic10: 1-78, Mic13: 1-118 | 0.61 |
| Homo-oligomer | | | | |
| 4, Mic60 | NA | Mic60$^{IMS}$ | Mic60: 451-597 | 0.18 |
| 2, Mic19 | NA | Mic19 | Mic19: 59-174 | 0.2 |
| 8, Mic10 | 7 PLM, 4 OLA, 4 MYR | Mic10 | Mic10: 1-78 | 0.35 |
| 4, Mic10 | 4 PLM, 4 OLA, 4 MYR | Mic10 | Mic10: 1-78 | 0.37 |
| 2, Mic10 | 4 PLM, 3 OLA, 3 MYR | Mic10 | Mic10: 1-78 | 0.72 |
| Hetero-oligomers | | | | |
| 1:1, Mic10-Mic13 | 4 PLM, 3 OLA, 3 MYR | Mic10, Mic13 | Mic10:1-78, Mic13:1-118 | 0.79 |
| 2:1, Mic10-Mic13 | 4 PLM, 4 OLA, 4 MYR | Mic10, Mic13 | Mic10:1-78, Mic13:1-118 | 0.57 |
| 1:1, Mic10-Mic13 | 4 PLM, 3 OLA, 3 MYR | Mic10$^{TM1, matrix,TM2}$, Mic13$^{TM}$ | Mic10: 13-60, Mic13: 8-23 | 0.81 |
| **2:1, Mic10-Mic13** | **4 PLM, 3 OLA, 3 MYR** | **Mic10$^{TM1, matrix,TM2}$, Mic13$^{TM}$** | **Mic10: 13-60, Mic13: 8-23** | **0.84** |
| 2:2, Mic10-Mic13 | 5 PLM, 5 OLA, 5 MYR | Mic10$^{TM1, matrix,TM2}$, Mic13$^{TM}$ | Mic10: 13-60, Mic13: 8-23 | 0.75 |
| 1:1, Mic60 -Mic13 | 4 PLM, 3 OLA, 3 MYR | Mic60$^{IMS}$, Mic13 | Mic60: 451-758, Mic13:1-118 | 0.71 |
| 1:1, Mic60-Mic13 | 4 PLM, 3 OLA, 3 MYR | Mic60$^{link, LBS1,LBS2, mitofilin}$, Mic13$^{central,C}$ | Mic60:598-758, Mic13:24-118 | 0.76 |
| 2:1, Mic60-Mic13 | 5 PLM, 5 OLA, 5 MYR | Mic60$^{link, LBS1,LBS2, mitofilin}$, Mic13$^{central,C}$ | Mic60:598-758, Mic13:24-118 | 0.41 |
| **1:1, Mic60-Mic13** | **4 PLM, 3 OLA, 3 MYR** | **Mic60$^{LBS1,LBS2,mitofilin}$, Mic13$^{central,C}$** | **Mic60: 627-758, Mic13:24-118** | **0.81** |
| 2:2, Mic60-Mic13 | 5 PLM, 5 OLA, 5 MYR | Mic60$^{LBS1,LBS2,mitofilin}$, Mic13 | Mic60: 627-758, Mic13:24-118 | 0.44 |

| | | | | |
|---|---|---|---|---|
| 1:1, Mic60-Mic10 | 4 PLM, 3 OLA, 3 MYR | Mic60$^{IMS}$, Mic10 | Mic60: 451-758, Mic10:1-78 | 0.79 |
| 1:1, Mic60-Mic10 | 4 PLM, 3 OLA, 3 MYR | Mic60$^{link, LBS1,LBS2,mitofilin}$, Mic10 | Mic60: 598-758, Mic10:1-78 | 0.8 |
| **1:1, Mic60-Mic10** | **4 PLM, 3 OLA, 3 MYR** | **Mic60$^{LBS1,LBS2,mitofilin}$, Mic10** | **Mic60: 627-758, Mic10:1-78** | **0.8** |
| 2:2, Mic60-Mic10 | 5 PLM, 5 OLA, 5 MYR | Mic60$^{LBS1,LBS2,mitofilin}$, Mic10 | Mic60: 627-758, Mic10:1-78 | 0.67 |
| 4:2, Mic60-Mic19 | 8 PLM, 7 OLA, 7 MYR | Mic60, Mic19 | Mic60: 1-758, Mic19: 1-227 | 0.32 |
| 4:2, Mic60-Mic19 | NA | Mic60$^{CC}$, Mic19$^{CC}$ | Mic60: 451-597, Mic19: 59-174 | 0.2 |
| 2:2, Mic60-Mic19 | 4 PLM, 3 OLA, 3 MYR | Mic60$^{LBS1,LBS2,mitofilin}$, Mic19$^{CHCH}$ | Mic60: 627-758, Mic19: 178-227 | 0.68 |
| 2:2:1, Mic60-Mic10-Mic13 | 6 PLM, 7 OLA, 7 MYR | Mic60$^{link, LBS1,LBS2, mitofilin}$, Mic10, Mic13 | Mic60: 598-758, Mic10: 1-78, Mic13: 1-118 | 0.49 |
| AlphaFold Database (Monomer predictions) | | | | |
| **AFDB ID** | **Lipids** | **Confident domains** | **Confident residue ranges** | **Average pLDDT of AFDB predictions** |
| **AF-Q5TGZ0-F1** | **NA** | **Mic10$^{N,TM1,TM2}$** | **Mic10: 2-36, 40-60** | **81.25** |
| **AF-Q5XKP0-F1** | **NA** | **Mic13$^{N,TM,central,C}$** | **Mic13: 1-68, 79-118** | **88** |
| **AF-Q9NX63-F1** | **NA** | **Mic19$^{CC}$** | **Mic19: 59-174** | **94.28** |
| **AF-Q16891-F1** | **NA** | **Mic60$^{link}$** | **Mic60: 598-626** | **88.88** |

**Table S3: Protein binding data used in modeling.** Protein-protein binding sites from biochemical experiments and AF3 predicted contacts used in integrative modeling are shown.

| Protein 1 | Region of protein 1 | Protein 2 | Region of protein 2 | Data used for modeling | Source of information |
|---|---|---|---|---|---|
| Mic10 | 1-78 | Mic13 | 81-85 | $Mic10^{1\text{-}78}$-$Mic13^{81\text{-}85}$ | Mutagenesis-western blot, native gel electrophoresis, co-IP [4] |
| Mic60 | 647 | Mic13 | 96 | $Mic60^{647}$-$Mic13^{96}$ | Interface residue pairs in contact from AF3 prediction |
| Mic60 | 652 | Mic13 | 96 | $Mic60^{652}$-$Mic13^{96}$ | |
| Mic60 | 656 | Mic13 | 99 | $Mic60^{656}$-$Mic13^{99}$ | |
| Mic60 | 657 | Mic13 | 96 | $Mic60^{657}$-$Mic13^{96}$ | |
| Mic60 | 660 | Mic13 | 95-96 | $Mic60^{660}$-$Mic13^{95\text{-}96}$ | |
| Mic60 | 660-661 | Mic13 | 92 | $Mic60^{660\text{-}661}$-$Mic13^{92}$ | |
| Mic60 | 663-664 | Mic13 | 92 | $Mic60^{663\text{-}664}$-$Mic13^{92}$ | |
| Mic60 | 664 | Mic13 | 88-89 | $Mic60^{664}$-$Mic13^{88\text{-}89}$ | |
| Mic60 | 667-668 | Mic13 | 88 | $Mic60^{667\text{-}668}$-$Mic13^{88}$ | |
| Mic60 | 668 | Mic13 | 88-89 | $Mic60^{668}$-$Mic13^{88\text{-}89}$ | |
| Mic60 | 668-669 | Mic13 | 85 | $Mic60^{668\text{-}669}$-$Mic13^{85}$ | |
| Mic60 | 669 | Mic13 | 81 | $Mic60^{669}$-$Mic13^{81}$ | |
| Mic60 | 669 | Mic13 | 84-85 | $Mic60^{669}$-$Mic13^{84\text{-}85}$ | |
| Mic60 | 688 | Mic13 | 85 | $Mic60^{688}$-$Mic13^{85}$ | |
| Mic60 | 641 | Mic10 | 3 | $Mic60^{641}$-$Mic10^{3}$ | |
| Mic60 | 641 | Mic10 | 6 | $Mic60^{641}$-$Mic10^{6}$ | |
| Mic60 | 653 | Mic10 | 62-63 | $Mic60^{653}$-$Mic10^{62\text{-}63}$ | |
| Mic60 | 655 | Mic10 | 62 | $Mic60^{655}$-$Mic10^{62}$ | |
| Mic60 | 658 | Mic10 | 62 | $Mic60^{658}$-$Mic10^{62}$ | |
| Mic60 | 720 | Mic10 | 3 | $Mic60^{723}$-$Mic10^{1\text{-}3}$ | |
| Mic60 | 720 | Mic10 | 6 | $Mic60^{723\text{-}724}$-$Mic10^{2\text{-}3}$ | |

**Table S4: Chemical crosslinks by mass spectrometry data used for integrative modeling.**

We obtained chemical crosslinks from XLinkDB and other studies.

| Species | Source of information | Crosslinker | Number of MICOS crosslinks |
|---|---|---|---|
| Mouse | [5] | BDP | 25 |
| Mouse | [6] | BDP | 30 |
| Human | [7] | BDP | 1 |
| Human | [8] | BDP | 3 |
| Human | [9] | BDP | 2 |
| Human | [10] | BDP | 4 |
| Human | [11] (HeLaInVivo) | BDP | 2 |
| Human | [11] (HeLaLysate) | BDP | 1 |
| Mouse | [12] | PIR | 64 |
| Human | [13] | PIR | 4 |
| Human | [14] | PIR | 5 |
| Human | [15] | DSSO | 13 |
| Human | [16] | DSSO | 42 |
| Human | [17] | DSSO | 11 |
| Human | [18] | DSSO | 47 |
| Human | [19] | BS3 | 81 |
| Human | [16] | DHSO | 6 |

**Table S5. Membrane restraints.** The domains of MICOS subunits are restrained based on their membrane topology. The membrane localization is obtained from co-IP, pull down assays, protein subcellular fractionation, and liposome sedimentation assays in human cells. The membrane restraint is implemented by computing the radial distance ($rad$) in the XY plane from the center $(0, 0, 0)$, $rad = \sqrt{x^2 + y^2}$, for a bead with coordinates $(x, y, z)$ (Fig. S1). Additionally, the Z-axial restraint applied along the cylindrical axis restrains the subunits to the upper rim of the cristae junction. The cristae are modeled as hollow cylinders with inner (r) and outer (R) radii.

| Restraint | Localization in cristae | Implementation: Harmonically restraining the beads | Residues restrained | Reference |
|---|---|---|---|---|
| Transmembrane restraint | Inner mitochondrial membrane | $R > rad > r$ | $Mic13^{8-23}$ | [4], [20] |
| | | | $Mic10^{13-36, 40-60}$ | [21], [22] |
| IMS localization restraint | Intermembrane space | $r > rad$ | $Mic60^{410-758}$ | [23] |
| | | | $Mic19^{1-227}$ | [24] |
| | | | $Mic13^{24-118}$ | [4], [20] |
| | | | $Mic10^{1-12, 61-78}$ | [21], [22] |
| Matrix localization restraint | Matrix | $rad > R$ | $Mic10^{37-39}$ | [22] |
| | | | $Mic13^{1-7}$ | [20] |
| Z axial restraint | Cristae junction | $0 < z < 50Å$ | $Mic60^{410-626}$, $Mic10^{1-78}$, $Mic13^{1-118}$ | NA |
| | Above the cristae junction, interacting with SAM/TOB complexes | $-30Å < z < 0$ | $Mic19^{1-14}$ | [24], [25] |

**Table S6. Fit to the chemical crosslinks.** We report the fit to crosslinks used in the modeling, as well as those not used in the modeling.

| Crosslink type, Species | Percentage satisfied (%) | Number of crosslinks Mic19-Mic19 violated/Total crosslinks |
|---|---|---|
| **Crosslinks used in modeling** | | |
| BDP/PIR, HUMAN | 100 | 0/13 |
| BDP/PIR, MOUSE | 95.8 | 4/96 |
| BS3, HUMAN | 95.4 | 1/22 |
| DHSO/DSSO, HUMAN | 88.8 | 8/72 |
| DSSO, MOUSE | 58.33 | 15/36 |
| **Crosslinks not used in modeling** | | |
| BDP/PIR, MOUSE | 90 | 3/31 |
| BS3, HUMAN | 100 | 0/7 |
| DHSO/DSSO, HUMAN | 100 | 0/24 |
| DHSO/DSSO, MOUSE | 0 | 2/2 |

**Table S7. Interface residue pairs between modeled MICOS subunits.** The novel protein-protein interfaces between the modeled subunits are summarized.

| Protein 1 | Residues 1 | Protein 2 | Residues 2 | Interacting protein domains | Same or different copies of the protein in intra-subunit contacts | Consistent with AF3 predictions |
|---|---|---|---|---|---|---|
| **Intra-subunit** | | | | | | |
| Mic10 | 61-70 | Mic10 | 5-12 | C-N | Same | |
| Mic10 | 37-39 | Mic10 | 37-39 | Matrix-matrix | Different | |
| | 41-51 | | 41-44 | TM2-TM2 | | |
| | 45-56 | | 45-56 | TM2-TM2 | | |
| | 57-59 | | 49-57 | TM2-TM2 | | |
| | 61-70 | | 61-70 | C-C | | |
| Mic19 | 31-50 | Mic19 | 31-50 | N-N | | |
| | 41-50 | | 51-58 | N-N | | |
| | 51-58 | | 51-58 | N-N | | |
| | 85 | | 84 | cc-cc | | |
| Mic60 | 620-623 | Mic60 | 747-748 | link-M | | |
| | 624-626 | | 744-749 | link-M | | |
| | 649-653 | | 724-735 | LBS2-M | | |
| | 649-653 | | 736-742 | LBS2-M | | |
| | 645-653 | | 647, 649-653 | LBS1,2-LBS1,2 | | |
| | 679-683 | | 646-647 | LBS2-LBS1 | | |
| | 649-653 | | 724-740 | LBS2-M | | |
| | 750-758 | | 619-626 | link-M | | |
| **Inter-subunit** | | | | | | |
| Mic60 | 648 | Mic13 | 89,92 | LBS1-C | | |
| | 664-668 | | 69-78, 80-97 | LBS2-central,C | | Yes |
| | 649-653 | | 88-101 | LBS2-C | | Yes |
| | 654-663 | | 87-105 | LBS2-C | | Yes |
| | 669-673 | | 79-92 | LBS2-C | | Yes |
| | 674-683 | | 79-89 | LBS2-C | | |
| | 679-683 | | 92 | LBS2-C | | |
| | 674-683 | | 79-89 | LBS2-C | | |
| | 686 | | 81-82, 84, 86 | M-C | | |

| | | | | | | |
|---|---|---|---|---|---|---|
| | 684-687, 690 | | 85-89 | M-C | | |
| | 690, 720 | | 81-82 | M-C | | |
| | 687 | | 81, 92 | M-C | | |
| | 705-710 | | 81-90 | M-C | | |
| | 711-722 | | 81-96 | M-C | | |
| | 714-718, 722 | | 97-98 | M-C | | |
| | 718-719 | | 80 | M-C | | |
| | 723 | | 84-85, 90 | M-C | | |
| | 723-726 | | 88-89, 91-96 | M-C | | |
| | 729 | | 95 | M-C | | |
| | 732, 736 | | 96 | M-C | | |
| | 735, 739 | | 89, 92-93, 96 | M-C | | |
| Mic10 | 15-18 | Mic13 | 16, 18-23 | TM1-TM | | |
| | 19-27 | | 12-23 | TM1-TM | | |
| | 19-23 | | 8-14 | TM1-TM | | |
| | 25-34 | | 8-11 | TM1-TM | | Yes |
| | 28-30 | | 18-20 | TM1-TM | | |
| | 32-35 | | 12-13, 16 | TM1-TM | | |
| | 35-36 | | 9-10 | TM1-TM | | |
| | 61-70 | | 24-31, 37, 40-41, 44, 47-48, 51, 55, 84-92 | C-central,C | | |
| Mic60 | 638-639 | Mic10 | 1-3,6 | LBS1-N | | |
| | 640-648 | | 1-6 | LBS1-N | | Yes |
| | 643-648 | | 7-10 | LBS1-N | | |
| | 646-648 | | 11-12 | LBS1-N | | |
| | 644-647 | | 61-70 | LBS1-C | | |
| | 649-653 | | 2-3, 5-6 | LBS2-N | | |
| | 679-683 | | 3-12 | LBS2-N | | |
| | 647-683 | | 61-70 | LBS2-C | | Yes |
| | 654-658 | | 71-78 | LBS2-C | | |
| | 687-689 | | 1-10 | M-N | | |
| | 689-692 | | 2-3 | M-N | | |

| | | | | | | |
|---|---|---|---|---|---|---|
| | 718-720 | | 3-4, 6, 11 | M-N | | |
| | 719-720 | | 5-12 | M-N | | Yes |
| | 721-727 | | 2-10 | M-N | | |
| | 723-728 | | 1-2, 5 | M-N | | |
| | 728-729 | | 2-3, 6 | M-N | | |
| | 730-731 | | 1-6 | M-N | | |
| | 732-735 | | 1-3 | M-N | | |
| | 684-686 | | 1-12, 61-70 | M-N,C | | |
| | 716-730 | | 61-70 | M-C | | |
| Mic19 | 209 | Mic13 | 97 | CHCH-C | | |
| Mic60 | 451, 454 | Mic19 | 220-227 | cc-CHCH | | |
| | 451-458 | | 180-181 | cc-CHCH | | |
| | 451-454 | | 209-217 | cc-CHCH | | |
| | 546-553 | | 222-227 | cc-CHCH | | |
| | 545-554 | | 178-180 | cc-CHCH | | |
| | 549-553 | | 181-186, 220-227 | cc-CHCH | | |
| | 550-551 | | 210-211, 213-218 | cc-CHCH | | |
| | 545, 548-550 | | 175-177 | cc-central | | |
| | 620-624 | | 175-180, 182-188 | link-central,CHCH | | |
| | 598-599 | | 175-186 | link-CHCH | | |
| | 600-603 | | 178-180 | link-CHCH | | |
| | 616-623 | | 183-184 | link-CHCH | | |
| | 625-626 | | 182-195 | link-CHCH | | |
| | 621-624 | | 217-227 | link-CHCH | | |
| | 623-626 | | 214-216 | link-CHCH | | |
| | 649-653 | | 200-209 | LBS2-CHCH | | |
| | 654-658 | | 199-203 | LBS2-CHCH | | |

**Table S8. Likely pathogenic mutations from ClinVar.** ClinVar variants of uncertain clinical significance were obtained and filtered based on pathogenicity from AlphaMissense [26], [27]. Mutations are likely pathogenic based on AlphaMissense are shown.

| ClinVar Variant ID | Protein | Mutation | AlphaMissense score | Protein-protein interface(s) harbouring the mutation |
|---|---|---|---|---|
| 3109529 | MIC60 | Glu476Ala | 0.8237 | - |
| 3109530 | MIC60 | Arg485Gln | 0.7008 | - |
| 3109531 | MIC60 | Val573Leu | 0.5974 | $Mic60^{CC}$-$Mic60^{mitofilin}$ |
| 2287116 | MIC60 | Leu612Phe | 0.7004 | - |
| 2598173 | MIC60 | Arg634Gly | 0.8398 | $Mic60^{LBS1}$-$Mic13^{C}$, $Mic60^{LBS1}$-$Mic60^{mitofilin}$ |
| 3528950 | MIC60 | Tyr658Cys | 0.8191 | $Mic60^{LBS2}$-$Mic13^{C}$, $Mic60^{LBS2}$-$Mic10^{C}$ |
| 3528954 | MIC60 | Glu704Gly | 0.5641 | $Mic60^{mitofilin}$-$Mic19^{CHCH}$ |
| 2621679 | MIC19 | Arg35Gln | 0.9517 | - |
| 3266812 | MIC19 | Glu109Lys | 0.7921 | $Mic19^{CC}$-$Mic19^{CC}$ |
| 3832734 | MIC19 | Thr159Asn | 0.8012 | - |
| 2532420 | MIC19 | Thr201Asn | 0.6547 | $Mic19^{CHCH}$-$Mic60^{CC}$ |
| 4002062 | MIC19 | Cys204Tyr | 0.9979 | $Mic19^{CHCH}$-$Mic60^{CC}$ |
| 3266814 | MIC19 | Gly226Glu | 0.6734 | $Mic19^{CHCH}$-$Mic60^{mitofilin}$ |
| 2603808 | MIC10 | Glu3Lys | 0.6142 | $Mic10^{N}$-$Mic60^{LBS1,LBS2}$ |
| 3395971 | MIC10 | Lys9Asn | 0.94 | $Mic10^{N}$-$Mic60^{LBS1,LBS2}$, $Mic10^{N}$-$Mic13^{C}$, $Mic10^{N}$-$Mic10^{C}$ |
| 3126296 | MIC10 | Ala15Val | 0.6909 | $Mic10^{TM1}$-$Mic13^{TM}$ |
| 3126297 | MIC10 | Asp16Gly | 0.9677 | $Mic10^{TM1}$-$Mic13^{TM}$ |
| 4054819 | MIC10 | Gly70Arg | 0.7452 | $Mic10^{C}$-$Mic60^{mitofilin}$, $Mic10^{C}$-$Mic13^{central}$, $Mic10^{C}$-$Mic10^{C}$ |

**Table S9: Protein binding data not used in modeling.** Protein-protein binding sites from biochemical experiments not used in integrative modeling are shown.

| Protein 1 | Region of protein 1 | Protein 2 | Region of protein 2 | Data used for modeling | Source of information | Reference |
|---|---|---|---|---|---|---|
| Mic10 | 1-78 | Mic13 | 15-19 (GXXXG motif) | $Mic10^{1-78}$-$Mic13^{15-19}$ | Mutagenesis-Western blot, native gel electrophoresis, Co-IP | [4] |
| Mic60 | 1-758 | Mic13 | 84-103 | $Mic60^{1-758}$-$Mic13^{84-103}$ | Co-IP | [4] |
| Mic60 | 1-758 | Mic19 | 172-221 | $Mic60^{410-758}$-$Mic19^{172-221}$ | Pull down, immunoblotting | [23] |
| Mic60 | 371-590 | Mic19 | 1-227 | $Mic60^{410-758}$-$Mic19^{1-227}$ | Pull down, western blotting | [28] |

**Table S10: Fit to crosslinks in Mic19$^{CC}$ parallel and antiparallel configurations.** We report the fit to intra-Mic19 crosslinks for parallel and anti-parallel Mic19 coiled-coil dimer structures. These structures were obtained from CCCP [28].

| **Dataset (violation length in Å)** | **Number of intra-Mic19 crosslinks mapping to Mic19$^{59-174}$** | **Number of crosslinks satisfied in the parallel Mic19$^{59-174}$ coiled-coil** | **Number of crosslinks satisfied in the anti-parallel Mic19$^{59-174}$ coiled-coil** |
|---|---|---|---|
| **Crosslinks used in modeling** | | | |
| **BDP/PIR, HUMAN (52)** | 0 | 0 | 0 |
| **BDP/PIR, MOUSE (52)** | 52 | 46 | 38 |
| **BS3, HUMAN (35)** | 12 | 11 | 10 |
| **DHSO/DSSO, HUMAN (30)** | 36 | 27 | 21 |
| **DSSO, MOUSE (30)** | 21 | 12 | 12 |
| **Crosslinks not used in modeling** | | | |
| **BDP/PIR, MOUSE (52)** | 15 | 14 | 9 |
| **BS3, HUMAN (35)** | 4 | 4 | 4 |
| **DHSO/DSSO, HUMAN (30)** | 8 | 7 | 6 |
| **DHSO/DSSO, MOUSE (30)** | 1 | 0 | 0 |
| **Total crosslinks** | **149** | **121** | **100** |

**Table S11. Sequences of synthetic peptides used in this study.**

| Peptide Name | Description | Sequence (N → C) |
|---|---|---|
| ctMic10$^{N}$ | ctMic10 N-region peptide with C-terminal FLAG tag | MSDSTTTPPRSVSRPVSEALLNEKWDRDYKDDDDK |
| ctMic10$^{C}$ | ctMic10 C-region peptide with N-terminal FLAG tag | DYKDDDDKSLKQATREFKKQQQQQQ |
| 1× FLAG | FLAG control peptide | DYKDDDDK |
| ctMic10$^{N_short}$ | Shortened ctMic10 N-region with C-terminal FLAG | MSDSTTTPPRSVSRPDYKDDDDK |
